# Multi-OMIC analysis of Huntington disease reveals a neuroprotective astrocyte state

**DOI:** 10.1101/2023.09.08.556867

**Authors:** Fahad Paryani, Ji-Sun Kwon, Chris W Ng, Nacoya Madden, Kenneth Ofori, Alice Tang, Hong Lu, Juncheng Li, Aayushi Mahajan, Shawn M. Davidson, Anna Basile, Caitlin McHugh, Jean Paul Vonsattel, Richard Hickman, Michael Zody, David E. Houseman, James E. Goldman, Andrew S. Yoo, Vilas Menon, Osama Al-Dalahmah

**Author notes:** Corresponding authors; Osama Al-Dalahmah - lead contact; Vilas Menon.

## Abstract

Huntington disease (HD) is an incurable neurodegenerative disease characterized by neuronal loss and astrogliosis. One hallmark of HD is the selective neuronal vulnerability of striatal medium spiny neurons. To date, the underlying mechanisms of this selective vulnerability have not been fully defined. Here, we employed a multi-omic approach including single nucleus RNAseq (snRNAseq), bulk RNAseq, lipidomics, *HTT* gene CAG repeat length measurements, and multiplexed immunofluorescence on post-mortem brain tissue from multiple brain regions of HD and control donors. We defined a signature of genes that is driven by CAG repeat length and found it enriched in astrocytic and microglial genes. Moreover, weighted gene correlation network analysis showed loss of connectivity of astrocytic and microglial modules in HD and identified modules that correlated with CAG-repeat length which further implicated inflammatory pathways and metabolism. We performed lipidomic analysis of HD and control brains and identified several lipid species that correlate with HD grade, including ceramides and very long chain fatty acids. Integration of lipidomics and bulk transcriptomics identified a consensus gene signature that correlates with HD grade and HD lipidomic abnormalities and implicated the unfolded protein response pathway. Because astrocytes are critical for brain lipid metabolism and play important roles in regulating inflammation, we analyzed our snRNAseq dataset with an emphasis on astrocyte pathology. We found two main astrocyte types that spanned multiple brain regions; these types correspond to protoplasmic astrocytes, and fibrous-like - CD44-positive, astrocytes. HD pathology was differentially associated with these cell types in a region-specific manner. One protoplasmic astrocyte cluster showed high expression of metallothionein genes, the depletion of this cluster positively correlated with the depletion of vulnerable medium spiny neurons in the caudate nucleus. We confirmed that metallothioneins were increased in cingulate HD astrocytes but were unchanged or even decreased in caudate astrocytes. We combined existing genome-wide association studies (GWAS) with a GWA study conducted on HD patients from the original Venezuelan cohort and identified a single-nucleotide polymorphism in the metallothionein gene locus associated with delayed age of onset. Functional studies found that metallothionein overexpressing astrocytes are better able to buffer glutamate and were neuroprotective of patient-derived directly reprogrammed HD MSNs as well as against rotenone-induced neuronal death *in vitro*. Finally, we found that metallothionein-overexpressing astrocytes increased the phagocytic activity of microglia *in vitro* and increased the expression of genes involved in fatty acid binding. Together, we identified an astrocytic phenotype that is regionally-enriched in less vulnerable brain regions that can be leveraged to protect neurons in HD.

## Introduction

Huntington disease (HD) is an incurable neurodegenerative disease characterized by selective loss of specific neuronal populations leading to motor, cognitive and psychiatric impairment^1–7^. Although the disease-causing mutation is present in all brain cells, the severity of the disease varies across brain regions^5, 6^. New evidence suggests that the CAG repeats in exon 1 of the Huntingtin gene (*HTT*) can expand during the lifetime of the organism, and in different cell types at different rates, which may contribute to disease progression^8–11^. Moreover, research implicates mutant *HTT* (mHTT) expression in cortical and striatal neurons as a necessary substrate for striatal neurodegeneration^12^. Furthermore, increased vulnerability of striatal neurons to deprivation of the cortical-derived, neuroprotective BDNF has shed light on striatal neuronal vulnerability in HD^13, 14^. However, despite recent advances in our understanding of regional heterogeneity of neurodegeneration in HD, there is much to be learned about how astrocytic heterogeneity contributes to this phenomenon.

In addition to the expression of *HTT* in neuronal cells, previous work has established the role of glial changes in HD. HD affects oligodendrocytes, microglia, and astrocytes. We previously described oligodendrocytic pathology at the single nucleus level in HD^15^. Astrocytes, the subject of this study, have long been noted to be “reactive”, as judged by their enlargement and the increase in markers such as GFAP, present particularly in the severely affected areas like the striatum (Vonsattel et al. 2008^16^ for review). Studies in human brain and in mouse models suggest a loss of function phenotype, including, for example, a decrease in the major glutamate transporter, EAAT2/GLT1 both in mouse striatum^17–19^, human striatum^20^, and in the cortex^20, 21^.

Astrocyte pathology is well-documented in HD. Studies of mHTT expressing astrocytes showed these cells can elicit and exacerbate HD pathology in murine models^22, 23^. mHTT can impair baseline astrocyte function^18^ and may elicit inflammation^24^. Also, downregulating Htt in astrocytes can slow down disease progression in murine HD models^25^. Moreover, human fetal-derived astrocytes with mHTT displayed several transcriptionally abnormalities, including low levels of metallothionein-3 (MT3) and genes involved in fatty acid synthesis. Functional deficiency of astrocytes can recapitulate specific motor and behavioral phenotypes in HD^26^. HD pathology extends from function to morphology. In fact, HD astrocytes are structurally abnormal^27, 28^, and astrocyte maturation defects have been described in HD iPSC models^28^. Thus, there is evidence for cell-autonomous astrocyte pathology that contributes to neuronal injury in HD.

The interaction between astrocytes and neurons is central to HD pathology. In fact, the response of astrocytes to injury in HD appears most robust when both neurons and astrocytes express the mutant protein in murine models, suggesting that a large component of astrocyte pathology in HD is “reactive” to neuronal injury in the setting of compromised basic astrocyte functions^29^. In addition, astrocytes with mHTT derived from human glial progenitor cells can recapitulate motor aspects of the HD phenotype when implanted into mice, while normal astrocytes can ameliorate HD pathology when implanted into HD mice^30^. Finally, genetically modified astrocytes can be leveraged to rescue HD symptoms^31^ and impart neuroprotection in HD^32–34^.

The goal of this study is to systematically map astrocyte pathology across different brain regions in human HD. Building on our previous results showing that mHTT protein aggregates in astrocytes, and that cingulate cortex astrocytes exhibit phenotypic heterogeneity even within the same region^21^, we generated multimodal omics data and performed systematic analyses to define astrocytic phenotypic heterogeneity in HD and correlate this molecular heterogeneity to disease severity. We combined bulk level RNAseq from 76 samples from 20 patients across different HD grades and 10 controls, as well as patients with juvenile-onset HD, with associated lipidomics from 27 cingulate cortex samples to extract disease severity signatures that correlate with CAG repeat length and HD grade. We also performed snRNAseq on samples from the severely affected caudate nucleus, and relatively less severely affected cingulate cortex and nucleus accumbens^35^, and analyzed astrocyte states and then correlated them to neuronal vulnerability across different brain regions. We found that protoplasmic astrocytes differed from fibrous-like CD44 positive astrocytes in their association with neurodegeneration and discovered further regional heterogeneity in protoplasmic astrocyte pathology across vulnerable and resilient brain regions. We discovered a gene signature characterized by increased metallothionein gene expression, which we posited is neuroprotective. We further analyzed existing GWAS data and identified a SNP in the metallothionein gene locus, a key hallmark of this putative neuroprotective signature, that is associated with delayed disease onset. We then performed validation and functional experiments and mapped metallothionein protein expression *in vivo* and confirmed the neuroprotective function of metallothionein-3 in astrocytes in vitro. Finally, we discovered a novel function of metallothionein-3 expressed in astrocytes on microglial function and gene expression. Together, our results pave the way for astrocyte-centric therapeutic strategies to treat HD.

## Results

### Transcriptomic analysis of multiple anatomic regions of HD brains identifies disease severity-associated gene signatures

The pathology of HD has been studied most in the caudate nucleus, one of the earliest and most severely affected brain regions. Transcriptional pathology has been described in the caudate nucleus and other brain regions including the frontal cortex, motor cortex, and cerebellum using bulk RNAseq^1, 6, 36, 37^, and in the caudate nucleus using snRNAseq^38, 39^. To define the transcriptional signatures of human HD that are dependent on disease severity, we performed transcriptional analyses of different brain regions including the severely affected caudate nucleus, and less severely involved cingulate cortex and nucleus accumbens. The methodology is illustrated in **Figure 1A**. We analyzed 76 brain samples from controls (n= 20 samples from 11 donors) and individuals with HD (n=56 samples from 24 donors, including 16 samples for Juvenile onset HD - **Supplementary Table-1**) using bulk RNA sequencing. We further performed lipidomic studies on a subset of these donors to validate pathologies described below, and single nucleus RNAseq (snRNAseq) to define pathology in glial cells (see below) - **Figure 1A**. Furthermore, we measured the CAG repeat lengths in the same HD samples and from the same tissue block we used for RNAseq (**Supplementary Table-1**). The CAG repeats ranged from 40-71 repeats. A general inspection of the dataset in the reduced dimensional space (tSNE) is provided in **Figure 1B** which shows samples color-coded by condition and anatomic region, and by sex and sequencing batch in **Figure S1A**. Overall, HD samples were enriched in the upper half of the tSNE projections (**Figure 1B**). There were numerous differentially expressed genes (DEGs), and we show a subset of these genes in **Figure 1C**. The gene expression heatmap demonstrates upregulation of several genes involved in control of gliogenesis/stemness such as *NES, EGFR, GLI1, PTCH1, YAP1, POU4F2, SMAD4,* and *REST*, and downregulation of several genes involved in oxidative phosphorylation and mitochondrial function, as well as several known HD medium spiny neuronal genes such as dopamine receptor genes *DRD1, DRD2*, striatal identity gene *PCP4* (**Figure 1C** and **supplementary table-2**). Next, we asked to what extent were the DEGs shared between brain regions (**Figure 1D** and **Supplementary Table-2)**. It was interesting to see that most DEG’s were region-specific and not shared with other brain regions. The overlap between DEGs among brain regions was most notable between the accumbens and caudate on the one hand - both striatal regions, and accumbens and cingulate on the other hand - both less severely involved in HD (**Figure 1D**). With this rich dataset, we determined the effect of CAG repeat length on gene expression using a multi-variate regression analysis taking into account sex, age, and anatomic region as co-variates. The results identified 1092 genes with either positive (672) or negative (420) regression weights (**Figure 1E** and **Supplementary Table-2**, we refer to these genes as “CAG-correlated”). The genes with most positive regression weights included astrocytic genes (*AQP4*, *CD44*, *LIFR*, *P2YR1*, *OSMR*, *SERPINA3*^40^ - also called Alpha1-antichemotrypsin) and microglial genes (*TLR3*, *VSIG4*, *IL13RA1*, and complement genes). We confirmed that *CD44*, a membrane protein expressed in astrocytes, was indeed increased in the HD caudate nucleus (**Supplementary figure 4A-B**). Next, we performed KEGG pathway enrichment analysis of the CAG-correlated genes and DEGs shared between two or three regions (**Figure 1F-G** and **Supplementary Table-2**). As expected, CAG-correlated genes were enriched in pathways related to splicing, protein processing, autophagy, neurodegeneration, and lysosomal function (**Figure 1F**). Interestingly, in both gene sets, KEGG pathways involved in inflammation, cancer-related pathways, and lipid metabolism were significantly enriched (**Figure 1F-G**). Moreover, weighted gene correlation network analysis identified gene modules correlated with CAG repeats; these modules were enriched in genes involved in DNA damage response and T-cell mediated inflammation, and loss of connectivity of key astrocytic and immune gene modules (**Figure S2** - see supplementary results). Together, these results point to a significant contribution of glia, namely astrocytes and microglia, in the pathology of HD.

**Figure 1.**
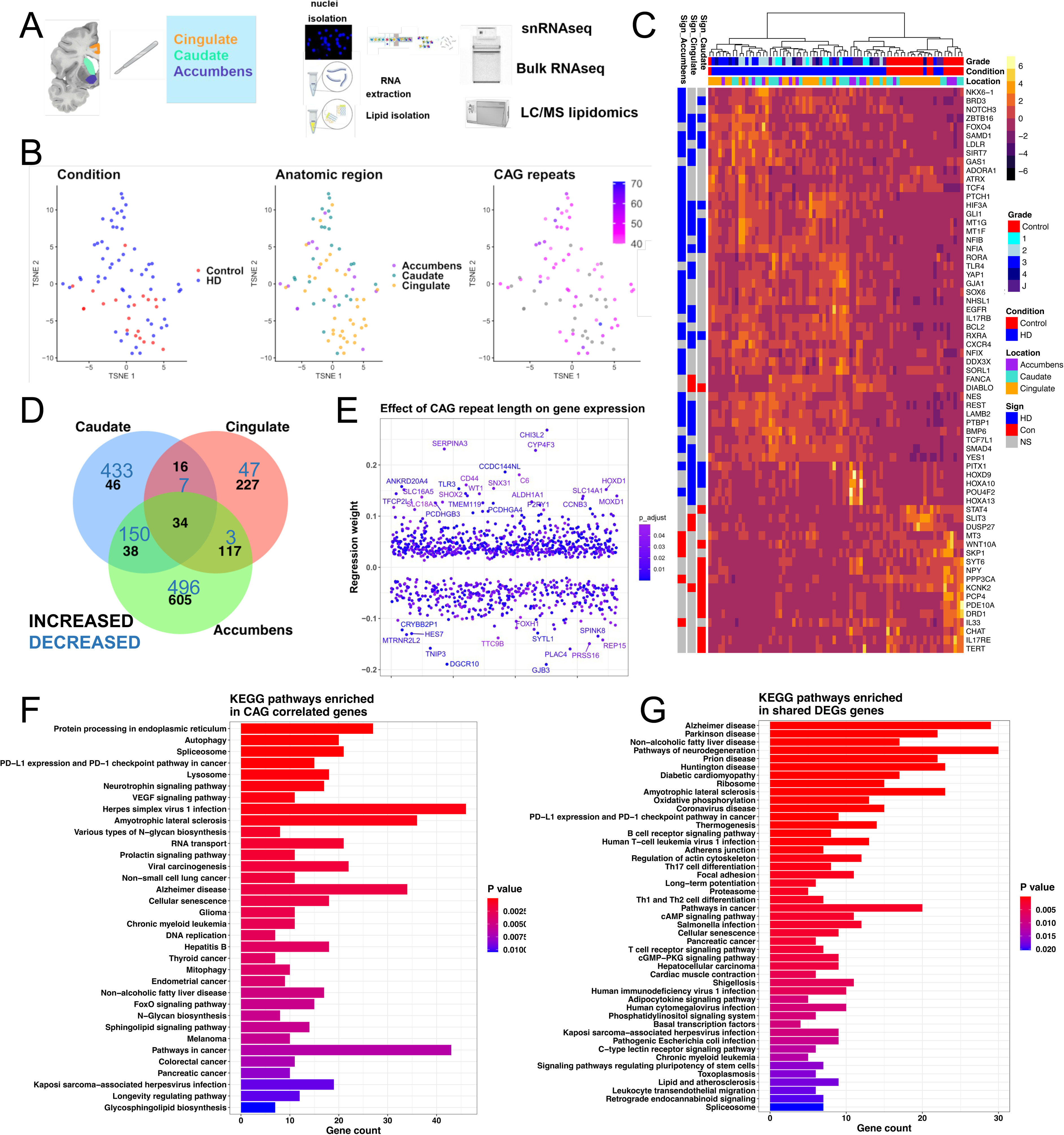
Transcriptomic analysis of HD identifies cross-regional and CAG-correlated gene signatures. **A**) Cartoon depicting experimental plan. **B**) t-distributed stochastic neighbor (tSNE) embedding of bulk RNAseq samples used in the study color-coded by condition (left), anatomic region (middle), and CAG repeat length (right). Control samples and ones with no available CAG repeat lengths are shown in grey. **C**) Heatmap of normalized gene expression showing a select subset of DEGs. The differentially expressed genes (DEGs - rows) are color-coded on the right by the direction of differential expression (Control -red vs HD - blue vs non significant (NS) - grey), and anatomic region where comparisons are significant (Sign_Acc: Accumbens, Sign_Caud: Caudate, Sign_Cing: Cingulate). The samples (Columns) are also color-coded by HD grade/Condition (Con: Control, HD1-4: 1-4, J: Juvenile onset HD). **D**) Venn diagram showing the overlap between DEGs - increased - black; and decreased - blue across the anatomic regions indicated. The genes consider for this analysis were considered significant if the adjusted p value was less than 0.05. **E**) Scatter plot showing genes with significant correlation with CAG repeat length. The correlation coefficient is shown on the y-axis, the order of genes on the x-axis is random. Color indicates adjusted p value. Genes with coefficients two standard deviations above the mean are indicated. **F-G**) EnrichR bar plots of KEGG pathways enriched in genes that positively or negatively correlate with CAG repeat length (**F**) or DEGs shared that are shared across two or three anatomic regions (increased and decreased - **G**).

### Integration of lipidomic analysis and transcriptomics implicates long-chain fatty acids and ceramides in HD neuropathology

Several studies have previously identified abnormalities in lipid species abundance in HD^41–47^. Whether certain lipid species vary with HD grade is still to be defined. To determine the relationship between HD grade and lipid species abundance, we performed lipidomic analysis of the cingulate cortex of 27 donors with different HD grades (**Supplementary Table-1**). We chose to examine this brain region because it is less severely involved in HD, and by examining different HD Vonsattel grades, we can determine what lipid species are correlated with disease progression. Regression analysis with grade as the explanatory variable and normalized lipid abundance as the response variable identified several lipid species that significantly varied over grade - including several long chain and very long chain fatty acids such as diacylglyceride (DG) and monoacylglycerides (MG), cholesterol, and lactosyl-ceramides (**Figure 2A**). Several of the very long chain monoglycerides were increased compared to control across different HD grades (**Figure S3A** **- Supplementary Table-4**). Given the differences between HD and controls in lipidomic signatures, we asked if we can predict HD grade from the lipidomics dataset. Thus, we performed sparse partial least square discriminant analysis (sPLSDA - **Figure 2B**) and found that HD grade can be accurately predicted from the loadings in the first four sPLSDA components - (**Figure S3B**). Our data analysis did not show significant effects of sex on lipid species expression as visualized in the sPLSDA space (**Figure S3C**).

**Figure 2.**
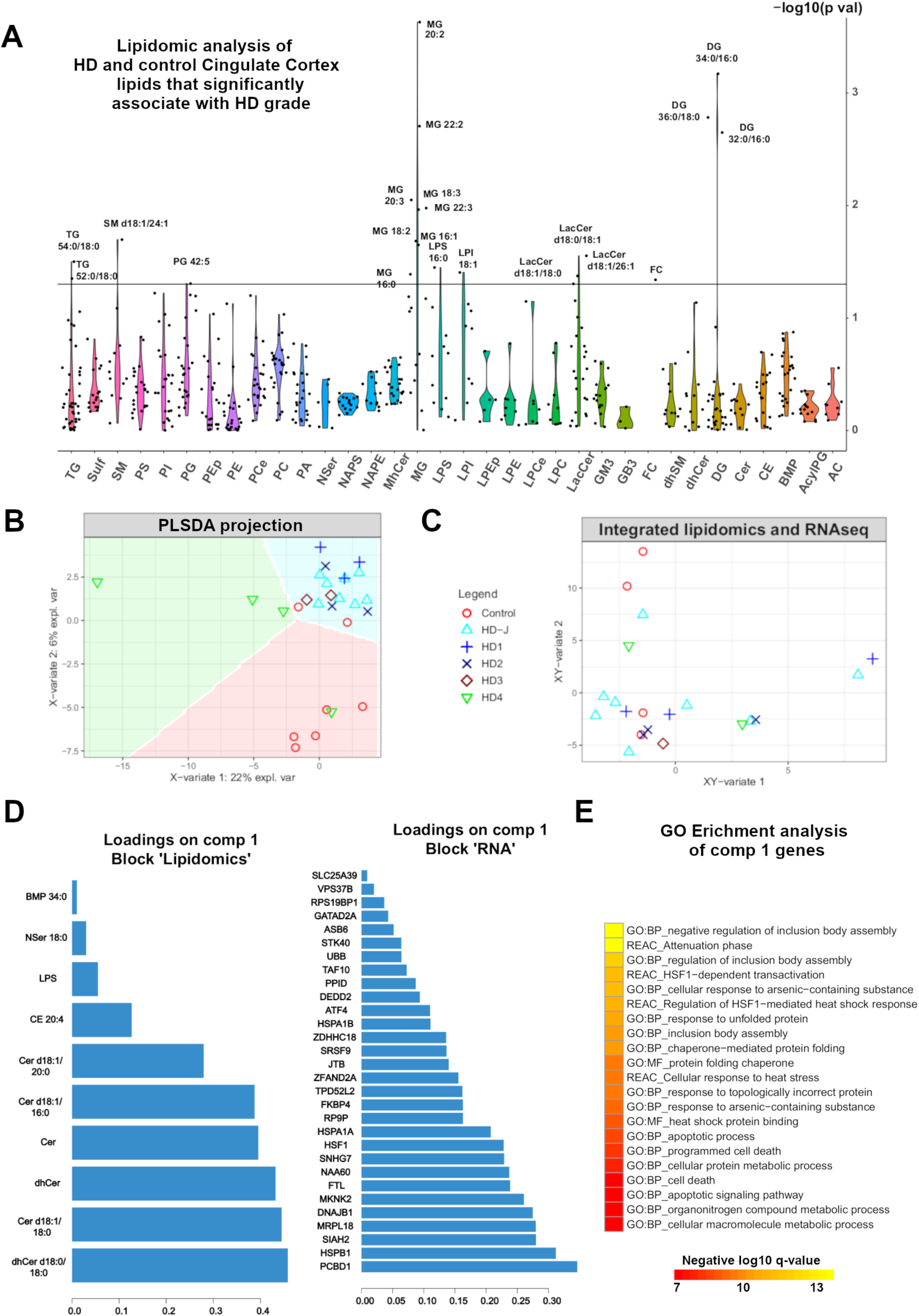
Lipidomic analysis of HD cingulate cortex. **A)** Violin plot of the -log10 of the p values of lipid species that significantly correlate with HD grade - see Figure S3A for details. **B)** Scatter plot showing the projection of lipidomics samples in the first two latent variables of the sparse-Partial least squares (sPLS) discriminant analysis model. The variance explained by each latent variable is indicated on the axes. The samples are color- and shape-coded by Condition/grade. The Condition can be predicted to a high degree of accuracy in the colored background regions - see also Figure S3B. **C)** Integration of lipidomics data and match bulk RNAseq data generated from the same samples using sPLS. The samples are color- and shape-coded as per B and projected in the combined integrated sPLS space. **D)** The loadings of the lipid species (left) and RNA transcripts (right) with strong positive correlation with the first sPLS latent variable that predicts grade are shown. **E**) Gene ontology enrichment analysis of component 1 genes. Negative log 10 of the adjusted p values are indicated.

Taking advantage of having RNAseq and lipidomics measurements from the same samples, we integrated a subset of 21 matched samples from both datasets using sparse-PLS to identify correlations between gene expression and altered lipid metabolism in HD. Projection of the integrated lipidomics and RNAseq data by HD grade and sex in the integrated space is shown in **figures 2C** and **S3C**, respectively. Unlike control samples, HD samples showed positive loadings in the integrated x-y variate-1 dimension (**Figure 2C**). Sex was correlated with the integrated x-y variate-2 dimension (**Figure S3C**). Several lipid species and RNA transcripts had positive loadings with the x-y variate-1, including lipids ceramides, dihydroceramides, and cholesterol esters, as well as several RNA transcripts (**Figure 2D**). Gene ontology enrichment analysis reveals the genes correlated with lipid of component 1, were enriched in ontologies related to cell death, apoptosis, inclusion body assembly, and response to unfolded protein (**Figure 2E**). The correlation network shows which and how RNA transcripts and lipids were correlated (**Figure S3D**). Altogether, our data indicate that lipid metabolism is altered in the cingulate cortex and link it to transcriptomic pathology, implicating the unfolded protein response as a central pathology that correlates with lipidomic pathology in HD.

### snRNAseq of HD brain reveals regional astrocytic heterogeneity

The results presented thus far are in accordance with our previous results implicating cingulate astrocytes in HD^21^, by pointing to a potential role of astrocytes in HD pathology; with numerous CAG-correlated genes and DEGs being astrocytic (e.g., *CD44*, *GFAP*), and changes in fatty acids abundance in HD - especially given the pivotal roles of astrocytes lipid metabolism^48^. We initially confirmed that CD44, a CAG-correlated gene, was increased in the caudate nucleus in HD at the protein level (**Figure S4A-B**). We also found that CD44 astrocytes were present in the pencil fibers, but only in HD, in the caudate nucleus parenchyma (**Figure S4A**). Together, these results implicate CD44 in HD pathology.

In order to characterize the heterogeneity of astrocytes along the axis of regional disease burden, we prepared snRNAseq samples from both control and HD brain samples from the nucleus accumbens, cingulate cortex, and caudate nucleus. We had recently reported the findings on oligodendrocytes and oligodendrocyte precursors in this dataset^15^. Here, we present the findings involving astrocytes, microglia, and neurons for the first time. Filtering and initial QC led to a total of 281,099 nuclei (**Supplementary Table-1**). Unsupervised clustering of these nuclei identified the major cell lineages as shown in the t-SNE plots space color-coded by lineage, region, and disease condition (**Figure 3A**). **Figure S5A** shows tSNE plots of the nuclei coded by donor, batch, and HD grade. **Figure 3B** shows the proportion of each lineage in each anatomic region, providing an overview of the dataset at hand. As expected, neurons predominated in the cingulate cortex, and ependymal cells were discovered only in the caudate nucleus. A subset of the canonical gene markers of cell type/lineage is shown in **Figure 3C**. Of all nuclei, 53,219 were astrocytes and after an additional rounds of QC to exclude doublets, ambiguous nuclei, or low quality cells, we recovered 45,101 high-quality astrocytes of which 7,556 were from control donors and 37,545 came from HD donors (**Figure 3D**, **Figure S5B**). The proportions of astrocytes in these three brain regions were roughly evenly distributed between cortical and striatal regions, allowing a comprehensive analysis of astrocyte regional diversity (**Figure 3B** and **Figure S5C**). In order to discover the underlying heterogeneity in the major astrocyte subtypes in our dataset and characterize their response to injury in HD, we classified astrocytes as protoplasmic - the most commonly studied astrocyte type, and fibrous-like, which include CD44 positive fibrous-like astrocytes that reside in the white matter, subependymal zone, perivascular regions, and subpial regions ^49 50^. In low-dimensional UMAP space, astrocytes varied along an axis with one end astrocytes expressed higher levels of *CD44*, *GFAP*, and *DCLK1*, and on the other end of the axis astrocytes expressed high levels of Wnt-inhibitory factor 1 (*WIF1*), Glutamine synthetase (*GLUL*), and *SLC1A2* (**Figure 3E**). To confirm our visual inspection, we subclustered the astrocytes and identified five clusters (**Figure S5D**), of which subcluster 0 harbored the majority of the CD44 expressing nuclei (**Figure S5E**). We designated that subcluster as fibrous-like (n= 18,700 nuclei), and the remaining clusters as protoplasmic (n = 26,401 nuclei). Because of the anatomic differences in the localization of these astrocyte types, we decided to re-analyze protoplasmic and fibrous-like astrocytes independently. We re-clustered both groups separately and found four clusters of fibrous-like astrocytes and seven clusters of protoplasmic astrocytes (**Figure 3F-G**). The gene markers of the fibrous-like and protoplasmic astrocytes are provided in **Figures 4A**, and **4E**, respectively (see also **Supplementary Table-5)**. Below, we present the fibrous-like and protoplasmic astrocytic clusters annotated by anatomic locale, clusters markers, and enrichment of specific informative gene signatures that we are interested: 1) a “putative” neuroprotective gene signature - which we defined based on our previous work to be enriched in metallothionein genes^51^ encoding proteins rich in cysteine amino acids known to confer protection against oxidative damage^52^, 2) CAG-correlated genes (see **Figure 1**), 3) genes associated with quiescent astrocytes^51^ - which are indicators of baseline astrocyte function including regulation of glutamate transport and glycolysis, and 4) genes correlated with the HD lipidomic signature - which are associated with unfolded protein response (see **Figure 2**). These genesets are provided in **Supplementary Table-2**. Finally, we determined the differentially expressed genes in protoplasmic and fibrous-like astrocytes per brain region examined (**Supplementary Table-6)**. Overall, as we show below, we were reassured to find that despite aggressive data integration, astrocytes clusters still reflected biological signal related to anatomic locale and adult-onset versus juvenile-onset disease.

**Figure 3.**
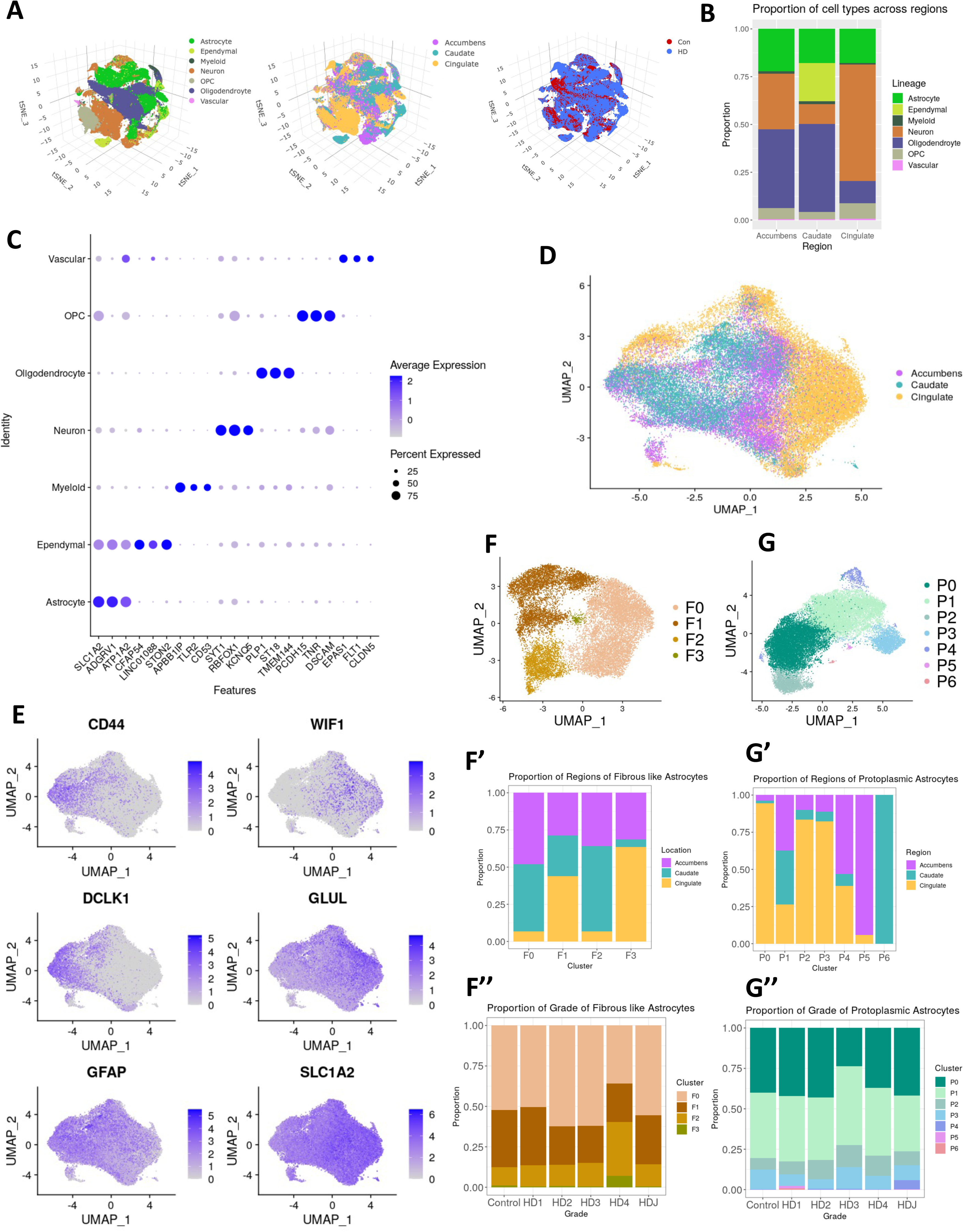
snRNAseq data analysis of HD and control astrocytes. **A**) tSNE projection of snRNAseq samples across all lineages (left), brain regions (middle), and condition (right). **B**) Stacked bar plot depicting the proportion (y-axis) of each cell lineage (color-coded) in different brain regions (x-axis). **C**) Dot plot showing select marker genes for each cell type/lineage. **D**) UMAP plot of the astrocytes snRNAseq profiles projected in isolation of other cell types, and color-coded by region. **E**) Feature plots of normalized gene expression projected in the UMAP embedding to highlight genes that differentiate fibrous-like (left) and protoplasmic astrocytes (right) - see also Figure S5. **F**) Sub-clusters of fibrous-like astrocytes (defined by highest expression of *CD44* - cluster 0 see Figures S5D-E) and a bar plot displaying the proportion of different brain regions in each cluster (**F’**) and the proportion of clusters in each HD grade/Condition (**F’’**). **G**) Sub-clusters of protoplasmic astrocytes (defined as all astrocytes except cluster 0 in Figure S5D) along with similar bar plots showcasing proportions of region (**G’**) in each cluster and proportion of cluster by HD grade/Condition (**G’’**).

**Figure 4.**
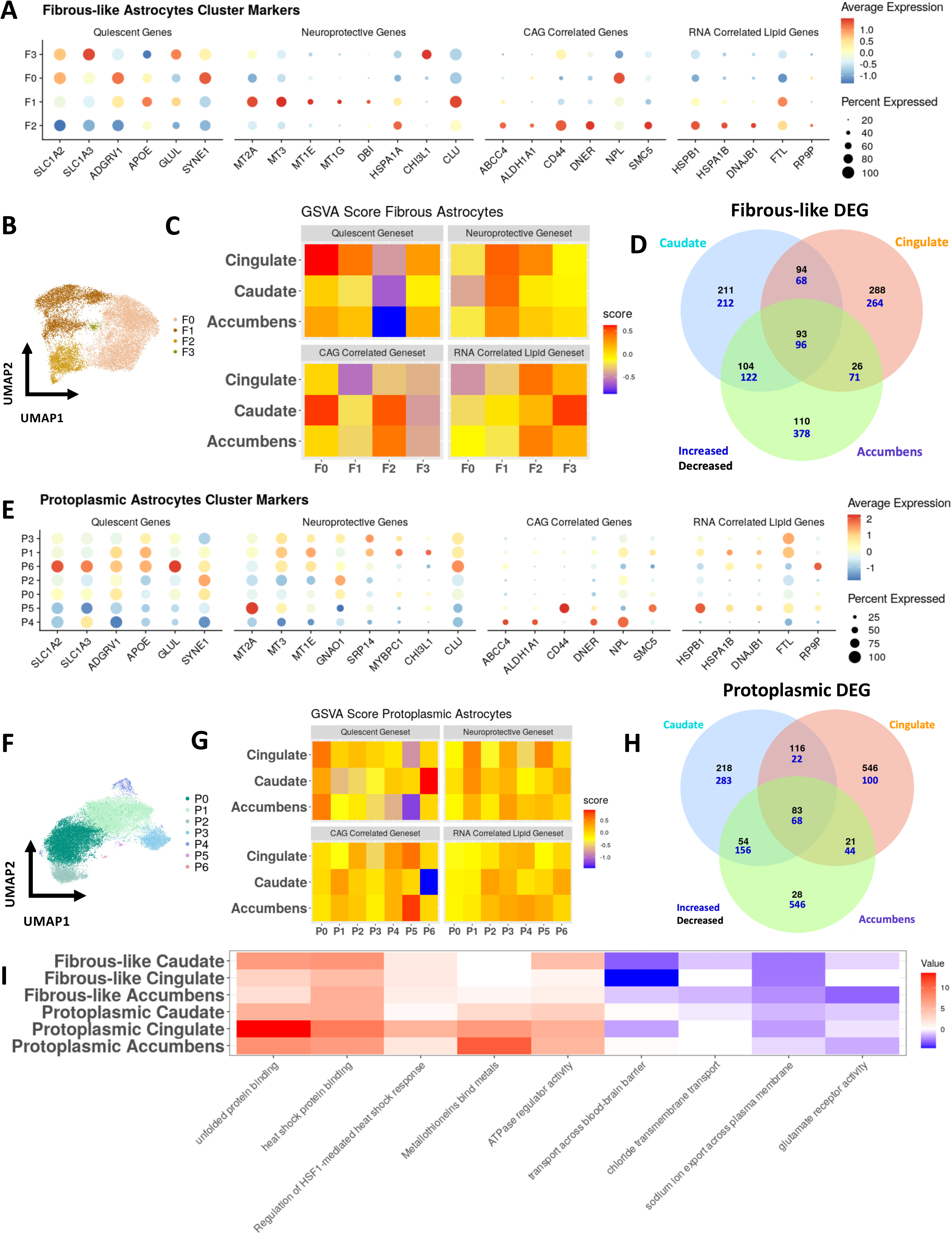
Astrocytes are regionally heterogeneous in HD. **A)** Dot plot displaying the expression of various genes from four different gene sets (Quiescent Genes = baseline astrocyte genes^51^, Neuroprotective Genes as predicted from our previous work^51^, CAG-correlated genes = genes with significant positive regression weights - see Figure 1E for more details, RNA-correlated Lipid Genes = set of genes that correlated with lipid abundance from Figure S3D. **B)** UMAP embedding of fibrous-like astrocytes. **C)** Heatmap of the average GSVA (gene set variation analysis) score within each cluster and brain region combination in fibrous-like astrocytes across the four gene sets described in A. **D)** Venn diagram analysis of DEGs in fibrous-across the three brain regions (increased: blue; decreased: black). **E)** Dot plot of same gene sets in **(A)** but for protoplasmic astrocytes. **F)** UMAP embedding of protoplasmic astrocytes. **G)** Similar GSVA analysis as **(C)** but for protoplasmic astrocytes. **H)** Similar DEG analysis as **(D)** but for protoplasmic astrocytes. **I)** Heatmap displaying the log10(p-value) of select GO terms (columns): red indicates terms significantly enriched in DEGs increased in HD, blue indicates GO terms enriched in DEGs significantly decreased in HD, and white indicates no significance. This is presented for fibrous-like and protoplasmic astrocytes (rows).

First, in the fibrous-like clusters, we found that majority of the cells are captured in clusters F0, F1, and F2 which contained 56%, 29%, and 14% of all fibrous-like astrocytes respectively. Cells from the accumbens and caudate were significantly enriched in clusters F0 and F2 (p values 1.69 x 10^-9^ and 1.57 x 10^-9^, respectively), whereas cingulate astrocytes were present mainly in clusters F1 and F3 as shown in the bar plot **Figure 3F’**. HD Vonsattel grades were represented in all clusters with no major deviations other than the observation that grade 4 samples contained a higher proportion of cluster F2 astrocytes (**Figure 3F’’**). Almost all cluster F3 astrocytes came from Juvenile-onset HD (**Figure 3F’’**). Clusters F0 and F3 show higher expression of glutamate transporters *SLC1A2* and *SLC1A3*, with the relatively small cluster F3 also showing expression of *GLUL* and *SYNE1*. Cluster F1 expressed metallothionein genes like *MT3, MT2A,* and *MT1E*, Clusterin (*CLU*), and APOE, while cluster F2 expressed heat-shock protein genes. Both of the aforementioned clusters F1 and F2 expressed relatively lower levels of glutamate transporters *SLC1A2* and *SLC1A3*. Multiple CAG-correlated gene and ones from the integrated lipidomics signatures were expressed highly in cluster F2. That said, cluster F3 is a relatively small cluster, and further interpretation is not pursued (**Figure 4A-B**).

Second, we set out to further understand the heterogeneity of both fibrous-like astrocytes. Initially, we interrogated fibrous-like astrocytic clusters using gene set variation analysis (GSVA) for the gene signatures we are interested in (see methods). We found several interesting cluster-specific and region-specific patterns (**Figure 4C**). The neuroprotective signature was enriched in cluster F1 in all regions, and in cluster F2 in the cingulate cortex. Also, the lipidomics associated geneset was most enriched in F2 astrocytes, which showed the highest expression of CD44, however, the pattern was less apparent in the caudate. The CAG-correlated genes were enriched in caudate cluster F0 astrocytes and in cluster F2 in the caudate and nucleus accumbens but not in the cingulate cortex. The quiescent geneset was enriched in the cingulate astrocytes in cluster F0, F1 and F3. Conversely, it was negatively enriched in cluster F2, also in a regionally-determined fashion; with the pattern being appreciated in the cingulate cortex and nucleus accumbens and less so in the caudate nucleus (**Figure 4C**). Together, these results show a strong regional influence on gene expression and likely functional states of fibrous-like astrocytes. Moreover, Cluster CD44-high cluster F2 displayed regional heterogeneity with expression of neuroprotective genes in the cortex, but CAG-correlated genes in the striatal regions.

As an additional analytic layer, we determined the differentially expressed genes (DEGs) which in fibrous-like astrocytes across all three brain regions. The numbers of DEGs are plotted in a venn diagram in **Figure 4D**. While there are shared DEG amongst the three brain regions, a large proportion of the DEG is uniquely attributed to the respective brain regions further highlighting the diversity of response to HD based on region. We discuss the DEGs below alongside DEGs from protoplasmic astrocytes to facilitate astrocyte type comparison.

Third, we considered protoplasmic astrocytes. For starters, cingulate astrocytes were most abundant in the most sizeable clusters P0, P2, and P3. Conversely, caudate and accumbens astrocytes were more represented in clusters P1 and P4. Clusters P5 and P6 were composed of cells from the nucleus accumbens and caudate nucleus, respectively (**Figure 3G’**). With the exception that most Juvenile-onset HD astrocytes were present in cluster P4 cells, no other discernable correlations were apparent between cluster and grade (**Figure 3G’’**). Examination of protoplasmic astrocytes cluster markers showed that while genes associated with baseline/quiescent astrocyte function were expressed in most protoplasmic astrocytes, they were highest in clusters P0 and P6. Clusters P1, P3 and P5 expressed elevated levels of metallothionein genes, and cluster P1 exhibited the highest levels of *CHI3L1,* and clusters P1, P3 and P6 expressed high levels of *CLU*. Clusters P4 and P5 showed expression of genes associated with the integrated-lipidomics signature and CAG-correlated genes (**Figure 4E-F**). That said, clusters P5 and P6 were small, and further interpretation is not pursued. Next, we interrogated protoplasmic astrocyte clusters using GSVA and DEGs. Using GSVA analysis of our genesets of interest (**Figure 4G**), cluster P0 was noticeably enriched in the quiescent gene signature, whereas clusters P1 and P3 were enriched in neuroprotective genes in all brain regions. Cluster P2 displayed regional heterogeneity and show high enrichment of CAG-correlated and lipidomics associated signatures in the striatal regions only. Cluster P4 astrocytes and caudate cluster P1 astrocytes showed highest scores for the CAG-correlated geneset scores, and cluster P3, in all brain regions examined, was enriched in the lipidomic gene signature (**Figure 4G**). The DEGs between control and HD protoplasmic astrocytes are shown in venn diagram across brain regions (**Figure 4H**), with the largest number of increased DEG being unique to the nucleus accumbens and the largest number of decreased DEG being found in the cingulate cortex.

Finally, to further analyze the DEGs in both protoplasmic and fibrous-like astrocytes across brain regions, we performed gene ontology (GO) enrichment analysis (**Figure 4I** **and Supplementary Table-6**. As expected, fibrous-like and protoplasmic astrocyte exhibited increased responses to unfolded protein and heat stress across all region and astrocyte types. Notwithstanding the similarities, there are significant differences in their response to injury that are type- and region-specific. For example, the ontology “metallothionein binding activity” is significantly increased in protoplasmic and not fibrous-like astrocytes - reflecting changes in levels of metallothionein genes. This pattern was more prominent in cingulate and accumbens astrocytes versus caudate astrocytes. Conversely, transportation across blood brain barriers is largely decreased in fibrous-like astrocyte with the largest effect taking place in the cingulate. Glutamate receptor activity was decreased most significantly in accumbens fibrous-like astrocytes. Altogether, protoplasmic and fibrous-like astrocytes exhibit diverse, region-specific, responses to injury in HD.

### Pseudotime analysis of gene variation reveals astrocytic gene programs

Our supervised analysis of astrocytic gene expression changes revealed that HD astrocytes exhibited diverse responses that vary between protoplasmic and fibrous-like astrocytes and across brain regions. To expand our insights into the dynamics of astrocytic response to injury, we took an orthogonal approach, agnostic to supervised gene signatures (neuroprotective, CAG correlated, and lipid correlated genes), and asked whether astrocytes vary across different gene programs. We reasoned that pseudotime analysis is a suitable approach because it can discover and describe the dynamics of branched trajectories of variation in gene expression in an unsupervised manner. In addition, it allows us to determine how HD and control astrocytes are differentially distributed along the trajectories of gene expression variation. We conceptualize these trajectories as gene programs that can increase our insight into astrocyte pathology in HD. Finally, this analysis allows us to computationally infer whether astrocytes can transition between transcriptional states, particularly from protoplasmic to fibrous-like.

To that end, we conducted pseudotime analysis on protoplasmic astrocytes in the cingulate cortex and caudate nucleus, and fibrous-like astrocytes in the caudate nucleus. We previously showed that CD44 protein expression is not significantly altered in HD in the cingulate cortex^21^, thus, we focused on fibrous-like astrocytes in the caudate nucleus, where we showed CD44 to be increased (**Figure S4A-B**). Before we proceeded, we confined our analysis to the largest clusters represented by multiple samples/HD grades, and therefore removed clusters P5 and P6 from protoplasmic astrocytes, cluster F3 from fibrous-like astrocytes due to their small size, and removed cells from HD4 donors because of low numbers.

To facilitate the analysis, we projected cingulate and caudate protoplasmic astrocytes, and caudate fibrous-like astrocytes in low-dimensional PHATE^53^ space (**Figure 5A and D**, respectively). We then examined the distributions of astrocytes in each trajectory by astrocytic cluster and HD Vonsattel grade (**Figure 5B-C** and **5E-F** for protoplasmic and fibrous-like astrocytes, respectively). Finally, by examining the genes differentially expressed in cells with high versus low loadings in each trajectory, we annotated each trajectory by meaningful ontologies and genesets using geneset enrichment analysis (**Figure 5G-H** and **Supplementary Table-7**).

**Figure 5.**
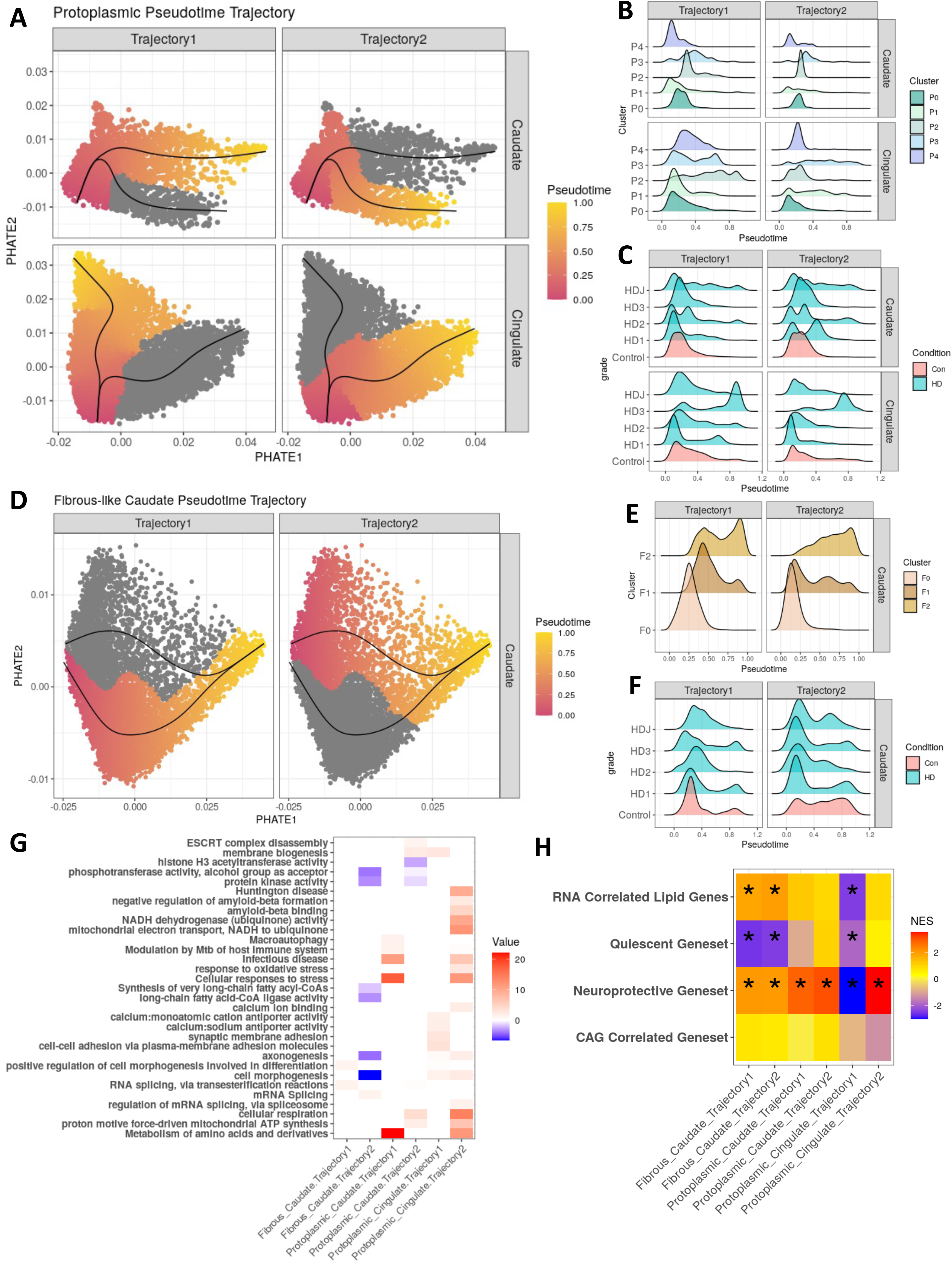
Pseudotime analysis reveals diverse astrocytic gene programs in HD. **A)** PHATE embeddings of protoplasmic astrocytes in the caudate nucleus and cingulate cortex along with pseudotime trajectories color-coded by pseudotime value. The principal graphs are indicated as black lines, and the cells are colored by their pseudotime values along each trajectory. In each plot, cells colored grey do not vary along the plotted trajectory. **B)** Histograms showing the pseudotime values of astrocytes as assigned by clusters (Figure 4F) in each trajectory and respective brain region. **C)** Same as B but showing astrocytes by HD Vonsattel grade. **D)** PHATE embeddings of fibrous-like astrocytes in the caudate nucleus along with pseudotime trajectories and color-coded by pseudotime values - same as A. **E)** Similar analysis as (**B**) but for fibrous-like astrocytes. **F)** Similar analysis as (**C**) but for fibrous-like astrocytes. **G)** Heatmap displaying the log10(p-value) of the GO terms enriched in the DEGs between the cells with highest vs lowest pseudotime values for each individual trajectory as implemented in Tradeseq. Red and blue colors indicate GO terms significantly enriched in DEGs increased and decreased along the trajectory, respectively. White color indicates no significance. **H)** Heatmap displaying the normalized enrichment score for GSEA analysis on our four gene sets discussed in (**Figure 4A**) with stars indicating if the enrichment is significant, using pre-ranked GO enrichment analysis.

First, we examined pseudotime trajectories in protoplasmic astrocytes, allowing us to discover the trajectories of gene expression along which astrocytes vary in both the cingulate and the caudate, as well as how astrocyte subclusters are distributed along the pseudotime axis (**Figure 5A & B**). In both trajectories, the pseudotime axes suggested variation from cluster P0 to P1/P3, with cluster P0 on one end in both regions examined, and cluster P3 on the other end. Only in trajectory 1 did protoplasmic cluster P2 display higher pseudotime values. Notably, having previously shown that protoplasmic cluster P2 was highly enriched with the CAG-correlated genesets, we contend that trajectory 1 represents a gene program characteristic of astrocytes with highest disease burden. Indeed, when projecting pseudotime values of protoplasmic astrocytes by HD grade (**Figure 5C**), we find increasingly higher pseudotime values in trajectory 1, however, that was not the case with Juvenile-onset HD in the cingulate cortex. Unlike the cingulate cortex, Juvenile-onset HD astrocytes adopted overall higher pseudotime values in the caudate. In the caudate nucleus, astrocytes appear to gain in pseudotime values in from HD grade 1 to 2, but then revert to control levels in HD3 in both trajectories, suggesting that astrocytes change their response to neurodegeneration as HD progress. These findings indicate that adult-onset and juvenile onset HD astrocytes behave differently in a region-specific manner.

Second, we found that Caudate protoplasmic gene program 2 was positively and negatively enriched for pathways involved in cellular respiration and H3 acetylation, respectively. Conversely, genes of cingulate protoplasmic astrocyte trajectory 1, which we posited was related to disease-burden, were negatively enriched in the neuroprotective geneset, quiescent astrocyte geneset, and even the lipidomics associated geneset (**Figure 5H**). Instead, they were enriched for ontologies associated with sodium-calcium antiporter activity, membrane biosynthesis, and cell adhesion (**Figure 5G**). Cingulate protoplasmic astrocyte trajectory 2, which was significantly enriched in our neuroprotective signature (**Figure 5H**), was also enriched for genes involved in pathways related to HD, response to stress, negative regulation of amyloid beta formation, and cellular respiration (**Figure 5G**). Interestingly, In the caudate, protoplasmic gene program 1 shared some pathways with Cingulate gene program 2, but was enriched for pathways related to immunological response like infectious disease and immune modulation - again highlighting regional diversity of HD protoplasmic astrocytes. Altogether, our findings highlight two modes of astrocytic response to injury in HD, a neuroprotective pathway, and another pathway associated with disease burden and increased expression of genes involved in membrane biogenesis and calcium signaling.

Third, when it comes to fibrous-like astrocytes, we realize that these astrocytes are heterogeneous and encompass subependymal, white matter, and perivascular astrocytes. However, we observed that astrocytes with protoplasmic morphology increase CD44 in HD in the caudate parenchyma, rather than the aforementioned compartments (**Figure S4A-B**). Thus, for this analysis, we were interested in the pathways that lead to “state transition” into a CD44 astrocyte. Therefore, we discovered trajectories that culminate in cells with the highest CD44 expression (cells with most positive PHATE1 embeddings in Figure 5C - see **Figure S6**). Two such trajectories became apparent in the caudate nucleus (**Figure 5D**). Projecting astrocytes by their respective clusters shows that cluster F0 cells in both trajectories had low pseudotime values, while cluster F1 and F2 cells showed higher values (**Figure 5E**). When projected by grade, it was interesting to see more astrocytes in HD showing lower pseudotime values in trajectory 2 (**Figure 5F**). This trajectory is interesting to us because several protoplasmic genes (*SLC1A2, C1orf61, ATP1A2,* and *ATP1B2*) decrease along this trajectory with increasing pseudotime values, nominating this trajectory as one along which “state transition” occurs.

Fourth, we found it difficult to annotate fibrous-like caudate astrocytes gene program 1 which showed no clear relationship to HD grade. Genes that increase along this trajectory where mostly enriched in ontologies related to RNA splicing and regulation of morphogenesis (**Figure 5G** and **Supplementary Table-7**). Conversely, genes that increased along caudate fibrous-like astrocyte gene program 2, the putative “state transition” gene program, displayed negative enrichment of pathways related to cell morphogenesis, and phosphorylation in kinases, and long chain fatty acid synthesis (**Figure 5G** and **Supplementary Table-7**). Given that fatty acid metabolism is a known protoplasmic or baseline astrocyte function^54, 55^, and in conjunction with upregulation of neuroprotective genes, and down-regulation of quiescent astrocytes genes (**Figure 5H**), we posit that this gene program may constitute a transition from protoplasmic to fibrous-like astrocytes. Compatible with this notion, we found that the expression of fatty acid transporters BLBP5 and BLBP7 was decreased in parenchymal caudate (but not cingulate) astrocytes in HD (**Figure S6A-D**). However, empirical evidence is needed to further support this notion.

As an additional attempt to understand the proposed “state transition” to CD44 positive astrocytes, we performed enrichment analysis for transcription factor binding sites in the gene that significantly increased or decreased along both trajectories in enrichR^56^ (**Figure S6E-F**) using the consensus CHEA and Encode transcription factors from ChIP-seq experiments dataset^57^. Interestingly, in both trajectories these transcription factors were enriched: *SUZ12, SMAD4* (which we found increased on the bulk level), *AR, NFE2L2, NANONG, ESR1*, and *GATA1*. Only in trajectory 2, one we associated with CD44 state transition, did we observe enrichment of *REST* (RE1 silencing transcription factor - also increased in HD samples on the bulk level), *POU5F1*, *TP53*, and *GATA2* transcription factors. Together, unbiased analysis revealed that astrocytes can adopt diverse gene programs in response to injury in HD that are regionally heterogeneous, some potentially neuroprotective, and others relating to possible state transitions from protoplasmic to fibrous-like. These trajectories appear to be regulated by similar transcription factors, but the transition to CD44 trajectory involves additional transcription factors such as REST.

Selective and differential neuronal vulnerability in HD across different brain regions Since specific neuronal types exhibit selective vulnerability to neurodegeneration in HD leading to type- and region-specific neuronal loss^58–60^, we turned our attention to neurons and examined changes in their abundance and gene expression across different brain regions. We are specifically interested in how neuronal pathology in HD is related to astrocytic phenotypes, as we discuss below. First, we started by sub-clustering striatal (n=19,155 nuclei) and cortical neurons (n=62,253 nuclei) separately. **Figure 6A** shows the major neuronal subtypes identified, which consisted of direct (d) and indirect (i) pathway spiny projections neurons (SPN) as well as several interneurons/GABAergic cells that expressed canonical markers, like LGR5, LINGO2, SST, NPY, and SYK. Select gene markers associated with these subtypes are shown in **Figure 6B**. **Figure 6C** shows cingulate cortex neuronal sub-clusters, with both GABAergic interneurons and projection/excitatory glutamatergic neurons. We identified layer (L) specific excitatory neurons from L2, L3, L4, L5, and L6 as well as GABAergic neurons by their expression of canonical gene markers, as shown in **Figure 6D**. The cluster markers are provided in **Supplementary Table-8.**

**Figure 6.**
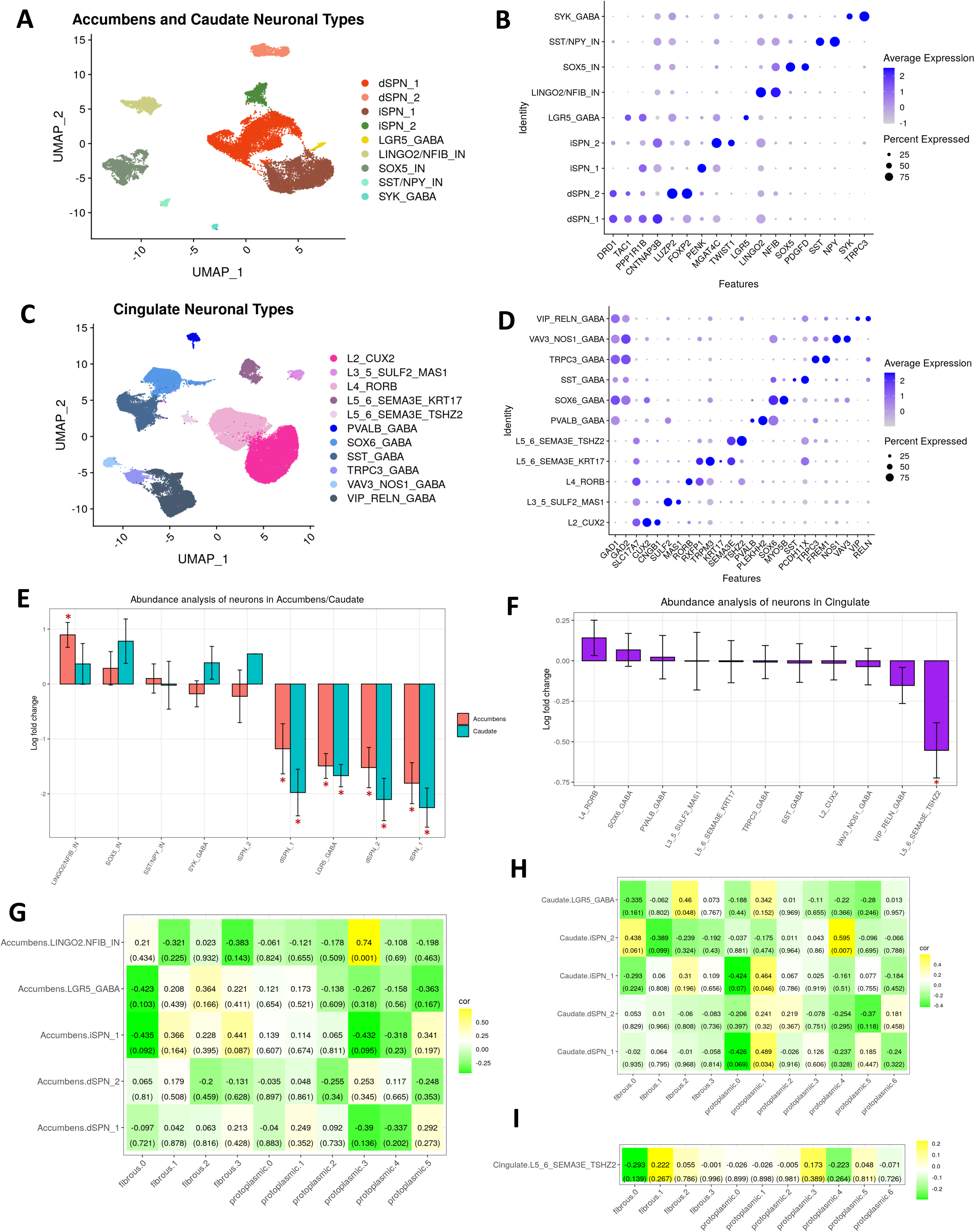
Differential abundance analysis of neurons in HD shows correlations to astrocytic states. **A**) UMAP plot of nucleus accumbens and caudate neuronal subtypes. **B**) Dot plot of gene markers that identify neuronal subtypes from accumbens and caudate cells in (**A**). **C**) UMAP plot of cingulate neuronal subtypes. **D**) Dot plot of gene markers for neuronal clusters from the cingulate (**C**). **E**) Differential abundance comparing the enrichment or depletion of neuronal subtypes in (**A**) in HD versus control. The logFC differences are shown on the y-axis. Starred plots indicate statistically significant differences. Error bars indicate SEM. **F**) Similar analysis as out but for cingulate cells from (**C**). **G**) Heatmap displaying the correlation of fibrous-like and protoplasmic sub-clusters, from Figures 3F-G, with proportions of neuronal clusters. The top value in each tile represents the Pearson correlation coefficient with its respective p-value below in parentheses. **H**) Similar analysis as (**G**) but for caudate neurons only from (**A**). **I**) Similar analysis as (**G**) but for cingulate neurons only from (**C**).

Next, we performed differential abundance analysis by comparing whether specific neuronal clusters were depleted in HD. In the caudate and accumbens, we found that dSPN_1, dSPN_2, iSPN_1, and LGR5_GABA were depleted in HD (**Figure 5E**). In the accumbens, LINGO2/NFIB_IN was relatively increased - likely reflective of apparent relative increase in proportion resulting from depletion of other neuronal types (**Figure 6E**). In the cingulate, only the L5_6_SEMA3E_TSHZ2 neurons were depleted (**Figure 6F**). Although SPN loss in the caudate has been described before^58, 59, 61, 62^, and in the nucleus accumbens^63, 64^, but to our knowledge, loss of LGR5+ interneurons has not been documented. Additional studies are needed to further characterize this phenotype.

Neuronal pathology at the snRNAseq in the striatum has been documented^39, 62^, and we provide an analysis of the DEGs in the striatal and cortical neurons in the supplementary results (**Figure S7** and **Supplementary Table 8**). We were interested in how patterns of neuronal loss correlate with astrocytic phenotypes. Thus, we quantified the correlations in abundance between neurons and the astrocyte sub-clusters to determine if certain populations of astrocyte were correlated with vulnerable neuronal clusters. The correlations between astrocyte clusters and neuronal clusters in the accumbens, caudate, and cingulate neurons are shown in **Figure 6G-I**. The heatmaps show the correlation coefficients and associated p-values in paratheses below. In the accumbens, the only significant correlation was between protoplasmic cluster P3 and LINGO2/NFIB inter-neuronal cluster, but due to small numbers of cells in this cluster, we refrain from making further remarks. In the caudate, there were significant positive correlations between fibrous-like cluster F2 and LGR5_GABA, and protoplasmic cluster P1 with dSPN_1 and iSPN_1, and between protoplasmic cluster P4 - which is harbored most Juvenile-onset HD protoplasmic astrocytes - and iSPN_2. In the cingulate cortex, there were no significant correlations (**Figure 6I**). The significant positive correlation between protoplasmic cluster P1 and the vulnerable MSN proportions which are depleted in the caudate is interesting because we hypothesized that protoplasmic cluster P1 astrocytes were neuroprotective based on our GSVA analysis. This led us to hypothesize that protoplasmic cluster P1 would be depleted in the caudate nucleus. And this is the subject of the validation studies below.

### Validation of astrocyte phenotypes in patient samples

We have shown that astrocytes showed an upregulation of a neuroprotective signature driven by metallothioneins, including MT3, in multiple brain regions (**Figure 4I**), we decided to validate it on the protein expression level. We were intrigued by this finding because it was most prominent in protoplasmic rather than fibrous-like astrocytes. We previously showed that cingulate astrocytes upregulate MT1 and MT2 genes in HD^21^. However, we have not quantified metallothionein protein expression in the severely affected caudate nucleus. So, we quantified MT1 and MT2A protein expression in caudate nucleus astrocytes (**Figure 7A**), and the found no significant differences between control and HD astrocytes (**Figure 7B**). We also examined the expression of MT3, which is also expressed in the putative neuroprotective astrocytes (**Figure 7C**) and found it to be decreased in parenchymal caudate astrocytes (**Figure 7D**). Conversely, in the cingulate cortex, we found that MT3, like other metallothioneins, to be increased (**Figure 7E-F**). This data is consistent with the finding of significant enrichment of metallothionein GO term in the cingulate more than the caudate (**Figure 4I**).

**Figure 7.**
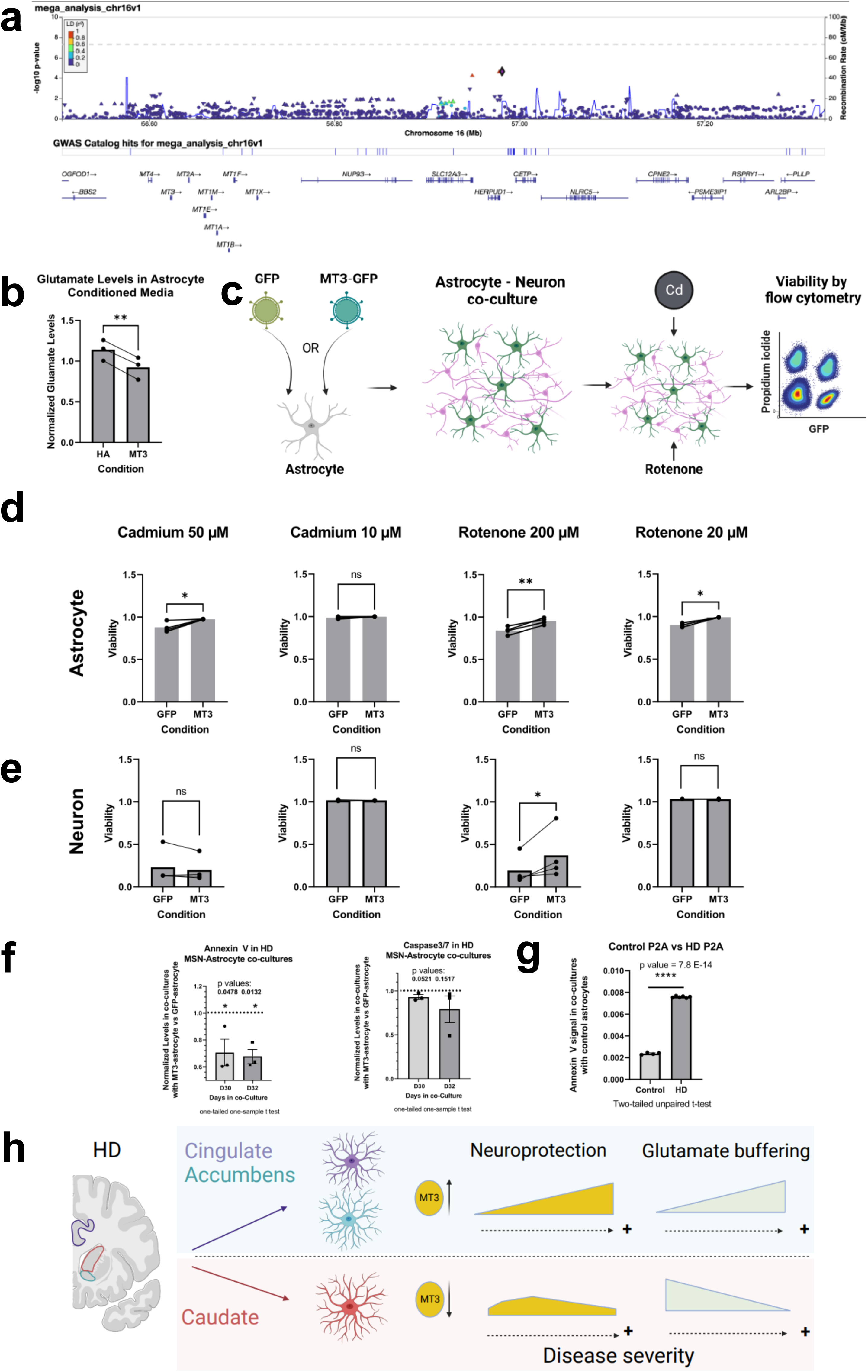
MT3 expression is Increased in the Cingulate and is Decreased in the Caudate of HD brains. Immunofluorescent images of the Caudate labeled for nuclei (DAPI-blue) and GFAP (green) to detect astrocytes (left), and MT (red-middle panel). A merged panel is shown on the right. Arrows indicate DAPI, GFAP and MT positive cells (MT positive astrocytes) and arrowheads indicate MT negative astrocytes. The antibody detects MT2A and MT1 proteins. Scale bar=50µm. Quantification of the percent of MT positive astrocytes in the Caudate. Unpaired one-tailed T-test with N=7 for control and HD. Data is shown as mean +/- SEM. P value= 0.3232. **C)** Immunofluorescent images of the Caudate labeled for nuclei (DAPI-blue) and GFAP (green) to detect astrocytes (left panel), and MT3 (red-middle panel). A merge of the three channels is shown on the right. Arrows indicate DAPI, GFAP and MT3 positive cells (MT3 positive astrocytes) and arrowheads indicate MT3 negative astrocytes. Scale bar=50µm. **D)** Quantification of the percent of MT3 positive astrocytes in the Caudate. Unpaired one-tailed T-test used with N=10 for control and 5 for HD. Data is shown as mean +/- SEM. P value= 0.0120. **E)** Same as D bur for the cingulate. **F)** Quantification of the percent of MT3 positive astrocytes in the Cingulate. Unpaired one-tailed T-test with N=8 for control and 6 for HD. Data is shown as mean +/- SEM. P value= 0.0405. For all IHC panels, control and HD images are shown on the top and bottom rows, respectively.

Next, we turned our attention to other markers of the putative neuroprotective protoplasmic Clusters P1 and P3. CLU and CHI3L1 (YKL40) were expressed in both clusters. CLU is a protein secreted from astrocytes and functions to ameliorate amyloid pathology in Alzheimer’s disease^65, 66^. CHI3L1 is also increased in Alzheimer’s disease astrocytes and is thought to be anti-inflammatory^67, 68^. We found it to be decreased in brain stem astrocytes in COVID-19 ^69^. It was interesting to observe both proteins to be expressed in the putative neuroprotective astrocytes. We thus quantified the expression of both proteins in the cingulate cortex and caudate nucleus. Quantification of CLU showed that it was decreased in the cingulate but not in caudate cortex HD astrocytes (**Figure S8A-D**). Conversely, CHI3L1 was increased in cingulate astrocytes and not caudate astrocytes (**Figure S8E-G**). Overall, these results show that even within the same cluster, astrocytes can show different patterns of protein expression. Thus, putative neuroprotective astrocytes in the cingulate are different from those in the caudate nucleus; they increase metallothioneins and CHI3L1 and decrease CLU, while caudate astrocytes of the same cluster did not exhibit altered expression of any of these proteins.

### GWAS analysis identifies a single nucleotide polymorphism in the MT gene locus associated with delayed disease onset

The age of onset of HD in patients is inversely correlated with the CAG length in the *HTT* gene. Once the effect of the CAG repeat length on age of onset is accounted for, the “residual” age of onset can be treated as a heritable trait. Using this trait, prior genome wide association studies identified several loci and genes as potential modifiers of the HD age of onset (see ^70^ for review). Here we directed our focus to the metallothionein gene cluster on chromosome 16, utilizing HD patient cohorts from both the GeM-HD consortium and Venezuelan Kindreds. We genotyped the Venezuelan Kindreds with a fine-mapping approach that combined whole genome sequencing and an Illumina CoreExome SNP array across 390 HD patients with corresponding clinical data. Our comprehensive mega-analysis, which combined the genotypic and clinical data from the Venezuelan patients with the GeM-HD consortium data^71^, identified three SNPs that are in linkage disequilibrium (rs3812963, rs74611520, and rs2518054) that were significantly associated with delayed age of onset in HD (**Figure 8A** and **supplementary table-7**). Further, we found that the two SNPs rs3812963 and rs74611520 act as an eQTLs and were associated with increased metallothionein levels in prefrontal cortex^72^ - see **supplementary table-7**.

**Figure 8.**
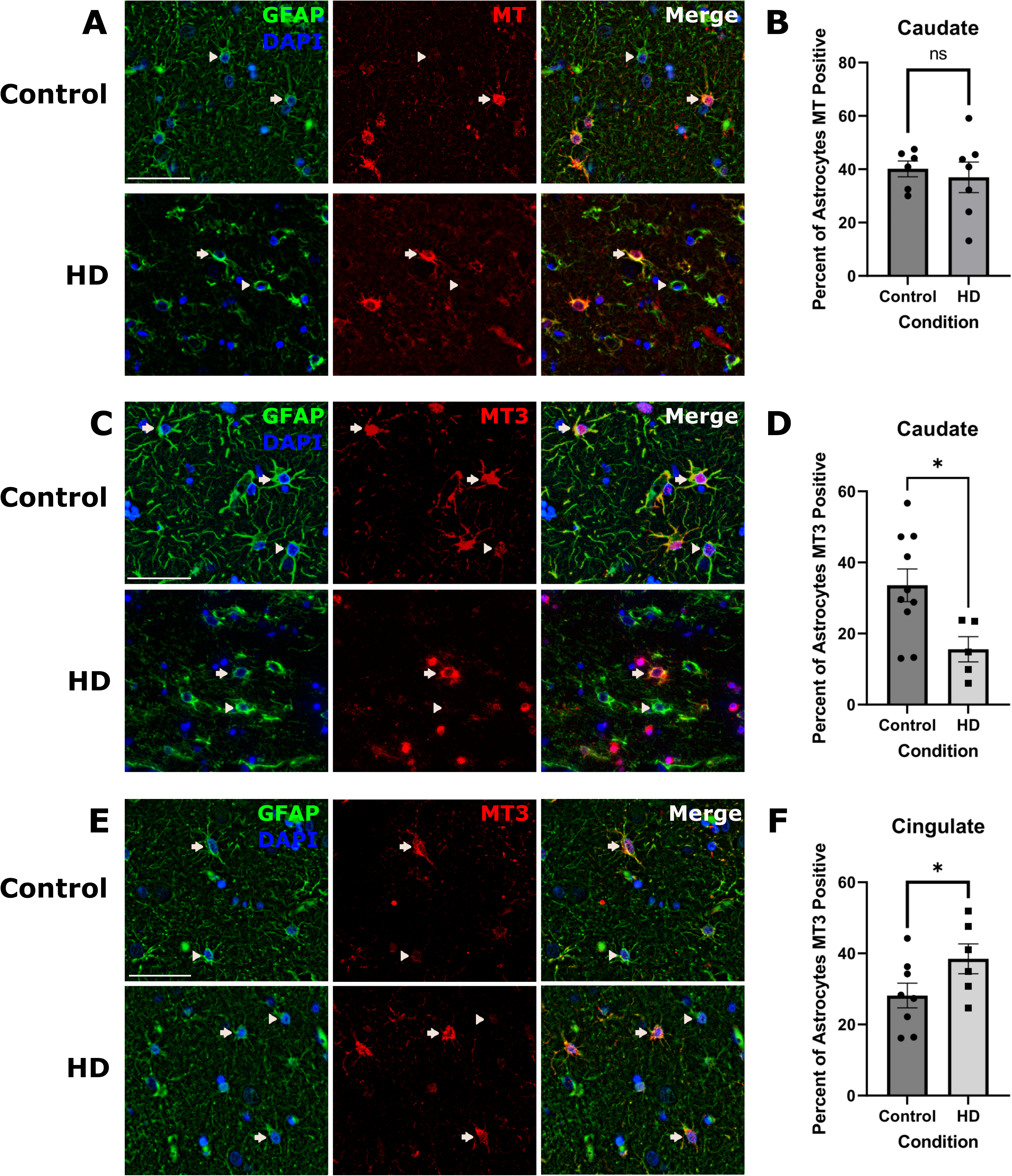
Metallothioneins are implicated in GWAS studies and are neuroprotective. **A)** GWA for HD residual age of onset identifies a significant signal within the metallothionein (mt) cluster of genes on chromosome 16. LocusZoom plot of the metallothionein (mt) gene cluster on chromosome 16 association with HD residual age of motor onset utilizing a linear mixed model. For each SNP in the association analysis, the log-transformed p-value of significance is graphed on the y-axis. SNPs are color-coded according to their correlation (r2) with the representative SNP rs74611520, as indicated by the diamond - the two neighboring SNPs that are in linkage disequilibrium with this SNP are indicated by the red triangles - rs2518054, rs3812963. This study combines data from the Venezuelan Kindreds and GeM-HD consortium of patients. **B**) Glutamate level measurement in conditioned media by control and MT3 astrocytes. **C)** A cartoon depiction of the design of the astrocyte-neuron co-culture viability experiment. **D)** Bar plots of the viability of control GFP vs MT3 overexpressing astrocytes when incubated in co-cultures with murine neurons under the indicated conditions. **E)** Bar plots of murine neuron viability when co-cultured with control GFP astrocytes GFP vs MT3 overexpressing astrocytes across the same set of conditions as in **D**. **F**) Expression of Annexin V and Caspase 3/7 in HD-derived directly reprogrammed MSNs co-cultured with control (GFP) versus MT3 astrocytes. The values are expressed as fold change from control. N=3 biological replicates. The p values are indicated. One-tailed one sample t-test. **G)** Example of Annexin V signal in control versus HD derived directly reprogrammed MSNs co-cultured with control astrocytes at day 30 demonstrating significant neurodegeneration in HD co-cultures evidenced by the increase in Annexin V signal. N= 4 and 6 technical replicates for control and HD, respectively, unpaired t-test, the p values are indicated. **H**) Cartoon illustration of astrocytic regional heterogeneity in HD and how that relates to neuroprotection.

### Metallothionein-3 is astroprotective and neuroprotective

We next sought out to define the functions of the putative neuroprotective signature. Thus, we modeled the neuroprotective astrocytes *in vitro* by overexpressing MT3 in human astrocytes (**Figure S9**) - which is increased in putative neuroprotective cluster P1 and P3. We selected MT3 over other metallothioneins because it was decreased in the HD caudate. We also overexpressed CLU, which is also expressed in clusters P1 and P3. MT3 overexpression selectively increased the expression of MT3 but not other metallothionein genes (**Figure S9A**). In CLU overexpressing astrocytes there was a modest upregulations of these CLU but no increase in metallothionein 1 genes (**Figure S9B-C**). Next, we asked if MT3 overexpressing astrocytes exhibited higher glutamate buffering capacity compared with control astrocytes. If so, that would suggest that MT3 astrocytes exhibited enhanced homeostatic functions. So, we measured glutamate levels in the conditioned medium of astrocytes cultured for 5-7 days. We found that MT3 astrocyte conditioned media contained lower levels of glutamate compared to control, suggesting MT3 astrocytes were better able to buffer glutamate **Figure 8B**. Consistent with this finding, MT3 astrocytes increased the expression of glutamate transporter *SLC1A2* and glutamine synthetase *GLUL* (**Figure S9D**). These results indicates that MT3 expression in astrocytes boosts glutamate buffering - a key astrocyte function can prevent neurotoxicity of glutamate.

We had predicted metallothioneins to be neuroprotective on the basis of reports showing MT1 and MT2 knockout mice exhibit increased vulnerability to neurodegeneration^73^. To our knowledge, the role of astrocytic MT3 was still untested. So, we tested whether MT3 astrocytes can protect murine neurons from heavy metal (Cadmium) or Rotenone-induced neurodegeneration as depicted in **Figure 8C**. We exposed astrocyte-neuronal co-cultures to Rotenone or Cadmium and measured cell viability to find that MT3 protected neurons against Rotenone-induced damage but not Cadmium induced damage (**Figure 8E**). Conversely, MT3 protected astrocytes from both types of damage (**Figure 8D**). Finally, we tested if MT3 astrocytes can neuroprotect directly reprogrammed HD patient-derived MSN^74^ (**Figure 8F**). Thus, we co-cultured control versus MT3 overexpressing astrocytes with HD-derived MSNs for 30 and 32 days, and measured the expression of cell death markers (Annexin V and Caspase3/7, n = 3 biological replicates). We found these markers to be decreased in the setting of MT3 co-culture (**Figure 8F**). HD-derived MSNs showed significant increase in Annexin V signal compared to control derived MSN (**Figure 8G**), supporting the utility of this model to test the effects of MT3 astrocytes on neurodegeneration^74^. Thus, our results overall showed that MT3 expression in astrocytes was not only neuroprotective in two models of neurodegeneration, but also astroprotective.

Interestingly, MT3 astrocytes also modulated microglial gene expression. In co-culture experiments of control or MT3-overexpressing astrocytes, we defined a microglial signature attributed to MT3 in astrocytes. Functionally, co-culture of MT3 overexpressing astrocytes with microglia led to increased microglial phagocytic activity. When examining microglial gene signatures in HD and control brain regions in our snRNAseq data, we found that the MT3 induced microglial gene signature we defined *in vitro* was enriched in HD microglia in the cingulate cortex - where astrocytes increased MT3 (see supplementary results).

## Discussion

In this work, we used transcriptomic and lipidomic analyses of different brain regions in HD to extract HD-related disease severity gene signatures. We found that these involve upregulation of glial genes including astrocytic and immune genes and downregulation of genes related to neuronal function. Lipidomic analysis of HD cortex revealed that long chain fatty acids were increased in HD. Integration of lipidomic and transcriptomics in HD implicated unfolded protein response genes as being correlated with HD lipidomic pathology. The unfolded protein response has been previously incriminated in HD pathology^75^. snRNAseq of protoplasmic and fibrous-like astrocytes allowed us to uncover astrocytic states that vary across brain regions and exhibit heterogeneous gene expression profiles. We identified a neuroprotective state of protoplasmic astrocytes which upregulated metallothioneins mainly in the cingulate cortex and nucleus accumbens, but was depleted in the caudate nucleus - where HD neurodegeneration was most severe. The depletion of the neuroprotective astrocytic state (protoplasmic cluster P1) was correlated with the depletion of vulnerable MSNs in the caudate nucleus. GWAS analysis incriminated a SNP, rs3812963, in the MT gene locus - which is associated with increased MT levels in the brain - to be associated with delayed disease age of onset. Functional experiments confirmed that metallothionein protein expression was neuroprotective, astroprotective, and induced increased expression of glutamate transporters, which increased glutamate buffering and lending further credence to the neuroprotective utility of metallothioneins in astrocytes. Finally, we found that genes induced in microglia due to astrocytic MT3 upregulation, which included genes related to fatty acid metabolism and lipid transport, could only be identified in microglia of the cingulate cortex, where astrocytes upregulated MTs, and the MT-primed microglia exhibited increased phagocytic capacity.

### Astrocytes in Different Areas Respond Differently in HD

Comparing snRNAseq profiles of astrocytes in the three variably affected brain regions in HD to control individuals, and across different grades of HD severity, revealed regional heterogeneity in astrocytic responses to HD. For example, caudate astrocytes failed to upregulate the putative neuroprotective proteins, metallothioneins 1 and 2, and even downregulated MT3. In contrast, cingulate astrocytes increased metallothioneins 1 and 2, as we previously reported^18^ and confirmed this in this study, and increased MT3 as well. Functional experiments revealed that metallothionein gene expression was indeed neuroprotective, as well as astroprotective. Metallothioneins are protective in cerebral ischemia, and other contexts of neuronal injury^76–79^. Most reports investigated MT1 and MT2 as neuroprotective molecules, but here, we found that MT3 is also neuroprotective. MT3 can regulate neuronal differentiation and promote survivability *in vitro*^80^, protect against seizure-induced cell death *in vivo*^81^, and is involved in the regulation of lysosomal functions and autophagy in astrocytes^82^. MT3 may be secreted via unknown non-canonical mechanisms^83^. The function of metallothioneins as chelators of heavy metals is well known^53, 84^, and given the increases in iron and copper in the HD brain^85^, the presence of metallothionines in astrocytes may be protective against oxidative stress. In this respect, the responses of cingulate vs. caudate astrocytes may be crucial for neuronal survival in these brain regions. Our finding that MT3 overexpression in astrocytes increased their ability to buffer glutamate is another potential mechanism by which MT3 confers neuroprotection - given the neurotoxic effects of excessive glutamate^86^. Future experiments will define the exact mechanism of neuroprotection conferred by MT3 expressing astrocytes.

We previously described a model for astrocytic responses to injury in the HD cortex, which was based on protoplasmic astrocytes^21^. In it we hypothesized that astrocytes which express quiescence genes at baseline respond to injury by upregulating neuroprotective genes and/or reactive genes like GFAP, then progressively decrease quiescence genes before they decrease neuroprotective genes. At end-stage, only disease-associated genes are present. Here, we see evidence to support this model in our expanded cohort of protoplasmic astrocytes across different brain regions. The addition of other anatomic regions as a dimension of progressive disease severity from cingulate to accumbens to caudate nucleus, makes a stronger case than the analysis of only the cingulate. Our model of astrocytic regional heterogeneity in response to HD is illustrated in **Figure 8H**.

### Different Types of Astrocytes Respond Differently

We separated astrocytes into two major categories, protoplasmic astrocytes and fibrous-like astrocytes, realizing that there is heterogeneity within each group. Fibrous-like astrocytes comprise a diverse group, including the interlaminar cortical astrocytes and white matter astrocytes in the normal human CNS (see our preprint^32^) as well as protoplasmic astrocytes that have diminished the expression of protoplasmic genes and acquired other genes, such as *CD44* and upregulated *GFAP*. Our neuropathologic examination of the HD caudate using IHC for CD44 suggested a transition from protoplasmic to CD44+ fibrous-like astrocytes. We described similar transition in different pathologies such as hypoxia and epilepsy^32^. To understand this phenomenon, we leveraged pseudotime analysis to define the gene programs that may regulate “state transition”. Interestingly, we identified pseudotime trajectory 2 (**Figure 5D**) which ended at a high *CD44* phenotype and progressed through a protoplasmic phenotype, defined by, the glutamate transporter *SLC1A2, C1orf61, ATP1B2,* and *ATP1A2* among others (**supplementary table 7**). This suggested that trajectory 2 represents a conversion of protoplasmic astrocytes to fibrous-like. This trajectory also shows a down-regulation of lipid-forming pathways, suggesting that the transition is characterized by a change in lipid synthesis and accumulation. Pseudotime trajectory 1 (**Figure 5D**), also ending at a high *CD44* phenotype, does not appear to go through a protoplasmic state, suggesting that these astrocytes may be endogenous CD44+ astrocytes, of which there are many in the normal caudate nucleus (see our preprint^32^). We did not try to distinguish among the different intrinsic CD44+ astrocytes, which will be the focus of a future study. The fibrous-like astrocytes in HD did not upregulate putative neuroprotective genes, including metallothioneins, as much as protoplasmic astrocytes. This suggests that the CD44+ astrocytes that are intrinsic to the normal CNS do not respond to HD in this way. The protoplasmic astrocytes that have become CD44+, more fibrous-like, also do not protect neurons in this way either.

To supplement the pseudotime analysis, we performed a ChEA analysis of TFs inferred from ChIP experiments^57^ and found a set of TFs enriched in the caudate fibrous-like astrocyte trajectories. Several of these are repressive TFs or components of repressive complexes that promote astrocyte differentiation over neuronal differentiation during CNS development, such as REST, which is increased by BMP signaling^87^. SUZ12 is a component of the polycomb repressor complex, which represses neuronal differentiation from neural stem cells in favor of astrocyte differentiation^88^ and is increased in murine HD astrocytes^28^. How these factors act in HD and other neurological disorders to cause changes in astrocyte states remains to be investigated.

### Associations between Astrocyte Responses and CAG Repeat length

Given that HD individuals carry variable numbers of CAG repeats, we were interested in associating various astrocyte clusters with the numbers of CAG repeats and found stronger correlations between clusters in the caudate than those in the cingulate cortex. This may not be surprising, since the caudate pathology is far more profound than the cortical pathology. The highest correlation appeared in the juvenile HD brains, which exhibited the largest CAG repeat lengths and expectedly showed high scores for CAG-correlated gene scores. Of interest, the neuroprotective protoplasmic cluster P1 displayed regional heterogeneity whereby it increased the CAG-correlated genes in striatal regions and not the cingulate cortex. This may be attributed to the differences in disease burden, but may also be due to potential inherent differences between human neocortical and striatal astrocytes. This is a subject of great interest to us and we will investigate it in a separate endeavor.

### Correlations between Astrocyte Pathology and Neuronal Loss

The known selective neuronal vulnerability in HD^58–60^ was recapitulated in our dataset. Our examination of neuronal populations in HD confirmed the loss the MSNs in the caudate, extended these results to the nucleus accumbens, and discovered the depletion of layer 5/6 SEMAS3E_TSHZ2 neurons in the cingulate cortex and a loss of LGR5+ interneurons in the caudate. We were particularly interested in any relationships between neuronal loss and astrocyte pathology, and so looked for correlations between astrocyte states and neuronal abundance. One of the significant positive correlations was between the neuroprotective protoplasmic cluster P1 and vulnerable MSNs. Since MSNs are depleted in the caudate, we reasoned that this neuroprotective astrocyte state might also be depleted, and indeed, our validation studies showed that there was a loss of MT3 in caudate astrocytes. It is possible that the selective neuronal vulnerability in the caudate nucleus is augmented by the lack of robust neuroprotective responses in caudate protoplasmic astrocytes, which we validated by IHC.

### Interactions between Astrocytes and Microglia

MT3 in astrocytes altered microglial gene expression and function, leading to increased phagocytic activity. In addition, genes induced in microglia due to astrocytic *MT3* upregulation, which included genes related to fatty acid metabolism and lipid transport, could only be identified in microglia of the cingulate cortex. Thus, astrocyte reactions in the cingulate may change microglial responses, which may be compensatory and contribute to the resilience of neurons in the cortex.

## Conclusion

In conclusion, we showed that astrocytes in HD exhibit state specific and regional heterogeneity in responses to HD pathology in human brain samples. We have defined gene signatures associated with CAG repeat length, lipidomic abnormalities, and neuroprotection. The latter was enriched in an astrocyte state that was depleted in the caudate nucleus and was correlated with the depletion of vulnerable striatal neurons. This neuroprotective gene signature was characterized by high levels of metallothionein gene expression. Analysis of HD GWAS studies identified a SNP associated with delayed disease onset in the MT gene locus, and functional experiments confirmed that MT expression in astrocytes was neuroprotective, and Astro-protective. Finally, we discovered a novel non-cell autonomous function of astrocytic MT3 on microglia, where it promoted microglial phagocytosis.

## Methods

### Bulk RNA sequencing

Total RNA was extracted from frozen brain tissue samples using an automated Qiagen platform (QiaSymphony). RIN values were measured on an Agilent Bioanalyzer. All samples had RIN values >= 7. The samples were processed using an Illumina® Stranded mRNA Prep kit and were sequenced on an Illumina NOVASEQ 6000 sequencer to produce stranded 100-base pair paired end libraries at 20 million read depth per sample. For RNAseq of microglial cultures, the samples were processed using an Illumina TruSeq Stranded total RNA kit and sequenced on an Illumina NextSeq sequencer at a read depth of 10 million reads per sample. Raw reads were aligned to the reference (GRCh38.92) using STAR^89^. Count matrices were generated from the BAM files using featureCount with default options. Differential gene expression analysis was performed in EdgeR using the glmQLFit method^90^. Batch was considered in the design formula. Gene set enrichment analysis was performed in enrichR^91^ and/or gprofiler2^92, 93^.

### CAG regression analysis

The gene expression count matrix (20276 genes) was filtered to remove genes with low expression using the “filterByExpr” function of the edgeR package (version 3.30.3), with default settings. 15413 genes were retained for subsequent analysis. The matrix was then transformed using the “limma::voom” function (version 3.44.3). Batch effects were visualized with principal component analysis and batch correction performed with the “limma::removeBatcheffects” function. To determine the how CAG repeat length affects expression of the retained genes in HD, we performed multiple linear regression with CAG as one of the predictors of the level of gene expression among HD samples. We assumed that level of gene expression would also be affected by age, region of the brain, and sex. A regression model was fit separately for each gene using the transformed gene abundance as the response variable, and CAG length, age, brain region from which a sample was procured, and sex, as predictor variables. Regression coefficients were estimated using least squares, and the p-values for the fit models were corrected for multiple hypothesis testing using the Benjamini-Hochberg method. Among the significant gene models (adjusted p-value < 0.01), genes which had a significant regression co-efficient estimate for CAG repeat length (p-value <0.01) were considered as the genes whose expressions were associated with CAG expansion - referred to as “CAG-correlated genes”. For enrichment analysis using EnrichR, we set an FDR threshold of 0.1 for select genes for the enrichment analysis.

### Lipidomics

Total lipids were extracted from frozen 40-70 mg human brain dissected as described above. Lipidomics profiling in mouse plasma and tissue samples was performed using Ultra Performance Liquid Chromatography-Tandem Mass Spectrometry (UPLC-MSMS). Lipid extracts were prepared from homogenized tissue samples using modified Bligh and Dyer method ^94^, spiked with appropriate internal standards, and analyzed on a platform comprising Agilent 1260 Infinity HPLC integrated to Agilent 6490A QQQ mass spectrometer controlled by Masshunter v 7.0 (Agilent Technologies, Santa Clara, CA). Glycerophospholipids and sphingolipids were separated with normal-phase HPLC as described before ^95^, with a few modifications. An Agilent Zorbax Rx-Sil column (2.1 x 100 mm, 1.8 µm) maintained at 25°C was used under the following conditions: mobile phase A (chloroform: methanol: ammonium hydroxide, 89.9:10:0.1, v/v) and mobile phase B (chloroform: methanol: water: ammonium hydroxide, 55:39:5.9:0.1, v/v); 95% A for 2 min, decreased linearly to 30% A over 18 min and further decreased to 25% A over 3 min, before returning to 95% over 2 min and held for 6 min. Separation of sterols and glycerolipids was carried out on a reverse phase Agilent Zorbax Eclipse XDB-C18 column (4.6 x 100 mm, 3.5um) using an isocratic mobile phase, chloroform, methanol, 0.1 M ammonium acetate (25:25:1) at a flow rate of 300 µl/min. Quantification of lipid species was accomplished using multiple reaction monitoring (MRM) transitions ^95, 96^ under both positive and negative ionization modes in conjunction with referencing of appropriate internal standards: PA 14:0/14:0, PC 14:0/14:0, PE 14:0/14:0, PG 15:0/15:0, PI 17:0/20:4, PS 14:0/14:0, BMP 14:0/14:0, APG 14:0/14:0, LPC 17:0, LPE 14:0, LPI 13:0, Cer d18:1/17:0, SM d18:1/12:0, dhSM d18:0/12:0, GalCer d18:1/12:0, GluCer d18:1/12:0, LacCer d18:1/12:0, D7-cholesterol, CE 17:0, MG 17:0, 4ME 16:0 diether DG, D5-TG 16:0/18:0/16:0 (Avanti Polar Lipids, Alabaster, AL). Lipid levels for each sample were calculated by summing up the total number of moles of all lipid species measured by all three LC-MS methodologies, and then normalizing that total to mol %. The final data are presented as mean mol % with error bars showing mean ± S.E. Statistical comparisons were done using a one-way ANOVA and Tukey’s test for post-hoc analysis. We had previously only presented the DAG results in our paper on oligodendrocytes^15^.

### WGCNA analysis

The “WGCNA” R package was used to identify modules of co-expressed and co-regulated genes, associated with various clinical and anatomic traits in HD^97^. Briefly, the gene expression count matrix of both control and HD samples was normalized using the voom function from limma^98^, batch corrected using combat^99^. To retain the most varying genes expressed across all samples, we determined the retained top 6183 genes with the highest median absolute deviation. A signed weighted gene co-expression network was constructed. Adjacencies of the retained genes were calculated using a soft thresholding power of 16 for control and 9 for HD networks. To minimize spurious associations, the adjacencies were transformed to a topological overlap matrix (TOM), and corresponding dissimilarity calculated. The genes were then clustered on the TOM-based dissimilarity. Modules of co-expressed genes were subsequently identified from the dendrogram of clustered genes using hybrid-tree cutting, with a deepsplit parameter set to TRUE and minimum module size of 75. Modules with similar expression profiles (cor > 0.75) were then merged. Significantly associated clinical and anatomic were then determined from the Pearson correlation between module eigengene expression and different traits.

### Dysregulated Networks of Co-expressed Genes in HD

To identify and characterize dysregulated gene expression networks in HD, we first built a reference network using control samples only following similar steps as above, and then evaluated the preservation of these modules in HD using the “modulePreservation” function in WGCNA with 100 permutations, as previously described^100^. Several module preservation statistics were generated, and used to determine the degree of preservation of each control module in HD.

### Integration of lipidomics with bulk RNAseq and sparse-PLS discriminant analysis

Integration of bulk RNAseq and lipidomics data was performed using the sparse-Partial least squares (spls) function in the R package mixOmics^101^. Both the normalized lipidomics and the log-normalized bulk RNAseq matrices were first filtered based on variance (by detecting outlier features with high variance using the isOutlier(da”a, nmads=2, type = “higher”) function from scater^102^. 5251 genes and 250 lipid species remained from 21 samples. The spls model was first tuned resulting in three components. The network was exported and visualized in cytoscape ^103^. Spls-discriminant analysis (splsda) was achieved after tuning the model using the following command tune.splsda(X = lipidomics_data, Y = Grade, ncomp = 4, test.keepX = c(1:10, seq(2”, 300” 10)), validation = “Mfold”, “olds = 5, di”t =””max.dist”, measure = “BER”, nrepeat = 10). The results x and y components were used in the final tuned model, which was used for discriminant analysis. ROC curve for sensitivity and true negative rates were generated using the auroc function from mixOmics.

### Clustering of various cell types from Huntington Disease samples

The main HD SingleCellExperiment object was created by aligning multiple datasets through the R package Seurat (version 4.06). Each sample’s cell type was identified using known markers and further corrected by region for biological consistency. The samples were log-normalized and scaled using Seurat’s NormalizeData function. Coordinates for the PCA were calculated using Seurat’s RunPCA function for a total of 30 principal components. The datasets were merged using Harmony (version 1.0) through Seurat’s RunHarmony function. All 30 principal components were used as inputs into RunHarmony. In addition, the integration corrected for three variables, donor, batch, and case numbers (some donors had more than one sample). UMAP coordinates were calculated with Seurat’s RunUMAP function using the Harmony coordinates as input into the dimensional reduction argument. Cluster assignment on the main object followed Seurat’s standard workflow with passing the object’s 30 Harmony coordinates through the FindNeighbors function followed by the FindClusters function with all default parameters. The coordinates of tSNE were produced using scater’s runTSNE function (version 1.22.0). Each tSNE plot was created using the first three coordinates through the package plotly (version 4.10.0).

### Sub-clustering of all Astrocyte samples

After clustering the main HD object which composed of various cell types, the astrocytes were subsetted into a separate Seurat object for reclustering. The samples were first log-normalized and scaled using Seurat’s NormalizeData function with default inputs. Afterwards, a PCA was performed using Seurat’s RunPCA function for a total of 30 principal components. All samples were then corrected for donor and batch through Seurat’s RunHarmony function. The inputs consisted of the first 30 principle components and the rest of the arguments were set to default settings. UMAP coordinates were calculated with Seurat’s RunUMAP function using the Harmony coordinates as input into the dimensional reduction argument. Cluster assignment on the main object followed Seurat’s standard workflow by passing the object’s 30 Harmony coordinates through the FindNeighbors function followed by the FindClusters function with all default parameters. Cluster assignments were evaluated again for any contamination by looking at standard markers for neuronal and glial cell types. Samples which consistently expressed certain cell markers were removed to ensure all samples were true astrocytes. To further sub-cluster astrocytes into fibrous and protoplasmic, we partitioned the astrocyte samples into those which expressed CD44 (fibrous-like) and does that did not (protoplasmic). These sub-types were then put into its own respective Seurat object and followed the same procedure of clustering as described above.

### Differential gene expression analysis - on the single cell level

An initial counts filter was applied to all DEG analysis for every single cell types. Depending on the cell type quality and whether pseudo bulking was applied, the counts filter criteria varied. The general methodology for determining the counts filter involved trimming down the total number of genes to test between 6000 10000. Significantly differential expressed genes between HD and Control were identified using limma (version 3.54.2) voom function with a regression formula to correct for Donor, batch, sex, and age. The analysis was followed with the eBayes function to apply statistics and finally extracting the top 1000 genes with an FDR of 5%. This methodology was applied to all cell types.

### Gene set variational analysis of various astrocytic states

The counts matrix for gene set enrichment analysis was prepared by pseudo-bulking the astrocyte snRNAseq data. The pseudo bulking procedure was performed by aggregating the counts from all donors within its respective UMAP cluster. All donor-cluster samples were then transferred into a Seurat object (version 4.06) and were normalized using Seurat’s NormalizeData function followed by the SCTransform function to correct for any covariates. All normalization functions used the default parameters. Gene set enrichment analysis was done using the GSVA package (version 1.44) where the input to the gsva function consisted of the pseudo-bulked matrix and the gene sets. The arguments of the function mx.diff was set to false and the method being “gsva”. From the output, we normalize the individual gene set scores into a z-score for each pseudo-bulk sample by applying the base R scale function. Finally, the gene set scores were averaged across location, grade, and cluster.

### Pseudotime analysis on astrocyte sub-populations

We created PHATE^53^ (version 1.0.4) embeddings for protoplasmic astrocytes from the cingulate and caudate, and fibrous-like astrocytes in the caudate individually. All raw counts matrix for each group were normalized using the sqrt function. We corrected for donors in the embedding by providing the phate function with a vector of all donors using fastmnn function, and used default parameters to create the final embedding. Pseudotime analysis was performed with slingshot^104^ (version 2.8.0) on the astrocytes in PHATE reduction. Initial clusters were defined using the Mclust package (version 6.0). Protoplasmic astrocytes starting points were defined by the highest enrichment of cluster P0 and fibrous-like astrocytes starting points were defined by the expression of CD44. Cluster annotations and PHATE embeddings were provided to the slingshot function with default parameters. Because pseudotime analysis is relative, setting multiple points of origin is not easily implemented in slingshot, and because we were interested in the different “routes” to a CD44 astrocytes, we set the origin point in the fibrous-like astrocytes to be the cell cluster with the highest CD44 expression and then inverted the pseudotime values by substracting them from 1. Downstream start vs end point differential expression was performed using the tradeSeq^105^ package (version 1.14.0).

### Differential abundance analysis

A matrix of cell type by donor for each brain region was created as input into the differential abundance analysis. Any donor that exhibited less 10 cell types from any particular cluster was discarded as an outlier. An otu table from the phyloseq package (version 1.42.0) was created for each matrix as required for running ANCOM-BC^106^ (version 2.02). The ancombc function was used to run the differential abundance analysis and with an input formula of “Age + Sex + Batch + Condition”. This insured our results were corrected for covariates. The prv_cut parameters was set to 0, neg_lb = TRUE, struc_zero = TRUE, conserve = FALSE, and all other parameters were set to the default setting. Log fold change values from the “Condition” column was used to compare whether specific cell types were enriched or depleted in HD. In addition, standard error values were extracted to place onto the final bar plot. Statistically significant cells were determined using the corrected p-value provided from ANCOM-BC.

### Correlation of astrocyte and neuron proportion analysis

Using the significantly differential abundant cells from the DA analysis, we correlated the proportions of significant cell types to the proportions of all astrocyte sub clusters for each brain region. The correlation was performed using the cor.test function from the stats package (version 3.6.2). All default parameters were used. The Pearson correlation and its respective p-value were extracted from the output of cor.test.

### Immunohistochemistry and multiplex immunofluorescence

5-7 micron-thick formalin sections of fixed and paraffin-embedded human brain tissue were processed on an automated Leica© Bond RXm autostainer according to the manufacturers’ instructions. For chromogenic DAB stains, generic IHC protocols were employed as per manufacturer protocols. Standard deparaffinization and rehydration steps preceded antigen retrieval in Leica ER2 (cat# AR9640) antigen retrieval buffer for 10-20 minutes according to manufacturer protocols. For multiplex immunofluorescence, one hour incubation in a blocking in 10% donkey serum containing PBS-based buffer preceded labeling with primary antibodies for 1 hour at room temperature. Three wash steps in Leica wash buffer (Ref#AR9590) preceded labeling with species appropriate Alexa fluor conjugated secondary antibodies (Invitrogen). After three washes, A DAPI containing mounting solution (Everbright TrueBlack Hardset Mounting Medium with DAPI Cat#23018) was used to label nuclei and quench autofluorescence prior to coverslipping. All steps were conducted at ambient temperature.

Alternatively, multiplex Immunofluorescence of paraffin-embedded blanks was done manually. First, the slides were rehydrated (xylene, 100% EtOH, 95% EtOH, 70% EtOH, 50% EtOH, ddH2O) and then washed in PBS-Tween based wash buffer. Slides were submerged in pre-heated 1xTrilogy for 20 minutes for antigen retrieval, cooled to room temperature, and washed in PBS-T. Samples were outlined with a hydrophobic marker to contain antibody solutions on the tissue. Slides were blocked with 1:10 dilution of blocking serum to AB diluent. Antibodies were diluted, applied, and slides were left to incubate 4°C overnight. On Day 2, slides were washed in PBS-T, species-appropriate secondary antibodies were diluted (1:500), applied, and slides were left to incubate at room temperature in the dark for 2 hours, and then washed in PBS-T. Slides were incubated with 0.1% Sudan Black B diluted in 70% EtOH for 30 minutes to quench the tissue’s autofluorescence, and then washed in PBS-T. Finally, slides were mounted with DAPI media to stain nuclei.

The following primary antibodies were used: Rabbit ALDH1L1 (1:100, EnCor, Cat#RPCA- ALDH1L1), Rabbit YKL-40 (1:250, Abcam, Cat#ab255297), Chicken GFAP (1:1000, Abcam, Cat#4674), Goat Clusterin (1:200, Thermo fisher, PA5-46931), Rabbit CD44 (1:100, Abcam, Cat#24illipore, Rabbit MT3 (1:100, millipore, Cat#HPA004011), FABP5 (1:100, Abcam cat#ab5674), FABP7 (1:1000 Sigma cat#HPA028825), GS (1:2500, BD Transduction cat#610518). Secondary antibodies conjugated to fluorophores: anti-mouse Alexa Fluor 488, 568, and 633, anti-rabbit Alexa Fluor 488, 594, anti-chicken Alexa Fluor 488 and 647, and anti-goat Alexa Fluor 488, 568, 633; all from goat or donkey (1:500, ThermoFisher Scientific, Eugene, OR).

### Imaging and image analysis

All brightfield images and immunofluorescent images were taken on the Leica Thunder imager DMi8. Images were acquired at 20x air or 40x oil immersion objectives using a Leica K5 camera or a Leica DMC5400 color camera. Leica biosystems LAS X software was used for image capture. Tiles covering the entire ION were taken and stitched. Leica Thunder instant computational clearing was used to remove out of focus light.

All observers were blind to experimental conditions. Tiff files were used for analysis in Qupath 0.4.2 Annotations delineating the dorsal caudate nucleus or the entirety of the cingulate cortex at the level of the caudate nucleus head were created. Next, to detect cells, we used the “cell detection” function (analysis tab), and the DAPI Channel was set as the Detection Channel. Next, we trained an object classified to classify the detections for the different channels. Training data was created from each image, for each channel, to mark the cells positive for a specific antigen.

One classifier per channel was trained by calling the “train object classifier” function under the analysis tab > classify with the following parameters: type=Random Trees, measurements = Cell: measurements = Cell: Channel X standard deviation, mean, max, and min measurements for the channel in question. To increase the accuracy of the classifier, additional training annotations were created on the image in question until the classification results matched the impression of the observer. A composite classifier was created from the individual trained classifiers using the “create composite classifier” function. Training images were included from all slides quantified, and classifiers were re-trained for each image separately as appropriate. For CD44 analysis, we created a pixel classifier to classify positive pixels. Training annotations were created for each image for positive and negative pixels. “Train pixel classifier” function was then called with the classifier type set to random trees, with a resolution of 2.60 µm/pixel, and selected all the features from only the channel in question.

### Statistics

For comparing astrocytes in HD and controls in multiplex IHC experiments, we used unpaired t-tests unless otherwise stated. All comparisons were two-tailed unless stated in the figure legends. For co-culture experiments, paired two tailed t-tests were used on the biological replicates, with each biological replicate representing the average of the technical replicates in one experiment. The n and p values are indicated in the respective figures.

### Culture and transduction of human astrocytes

Human astrocytes (Sciencell© cat#1800) were cultured in Astrocyte culture medium (Sciencell © cat#1801) according to vendor’s protocols on poly L-Lysine coated cell culture plates. The following Lentiviruses were obtained from VectorBuilder ™; Lentivirus transduction for mGFP (pLenti-C-mGFP-P2A-Puro - Origene Cat# RC211875L4V), CLU_mGFP (CLU(mGFP-tagged)- human clusterin(CLU) - Origene Cat# PS100093V), and MT3 (pLV[Exp]-EGFP:T2A:Puro- EF1A>hMT3 VectorBuilder cat# VB900003-8937eud).

### Real-time quantitative PCR

Total RNA was extracted from brain specimens using RNAeasy minikit (Qiagene©). RNA concentration and purity were determined using NanoDrop (Thermo Scientific™, MA). RNA was converted to cDNA using High-capacity RNA-to-cDNA kit (Thermo Fisher Scientific, Applied Biosystems™, MA). The following Taqman assays were used (GLUL cat #4453320, SLC1A2 cat# 44488952 and GADPH cat#4448484). The reaction volumes were 10-15 µl per reaction. TaqMan™ Multiplex Master Mix (Thermo Fisher Scientific cat# 4461881) was used. All reactions included 2-5 ng of cDNA. Thermal cycling parameters were conducted per manufacturer’s standard recommendations. The qPCR plates were read on a QuantStudio™ 5 Real-time PCR system (Thermo Fisher Scientific, Applied Biosystems™, MA). The reactions were done in triplicates. Relative gene expression was calculated using the delta delta Ct method with GAPDH as a reference gene. N = three biological replicates.

### Genome-wide genotyping and imputation of Venezuelan patients, and data acquisition from GeM-HD consortium

DNA was extracted from EBV-transformed lymphoblastoid cell lines established from blood samples of the Venezuelan individuals using the Qiagen Blood and Tissue DNeasy Kit. Genotyping on these DNA samples was performed at the New York Genome Center on a HiScan Illumina machine with the Illumina CoreExome Array. The genotypes were then imputed using the IMPUTE2 software package and the publicly available 1000 Genomes reference panel, along with whole genome sequences from select Venezuelan patient samples. For the GeM-HD consortium patients^107^, genotyping array data was downloaded from dbGaP (Study Accession: phs000222.v4.p2). The IMPUTE2 software package was used to impute these downloaded genotypes, guided by the 1000 Genomes reference panel.

### Genome-wide association study of residual age of onset

For the comprehensive mega-analysis, the genotypic and clinical data from the Venezuelan patients were combined with the data from the GeM-HD consortium. A quantitative genome-wide association (GWA) test was carried out using the GENESIS software package. A linear mixed model regression was implemented using the residual age at motor onset as a phenotype, allowing for adjustment of empirical pairwise relatedness, as well as other covariates such as sex and population structure. The residual age of onset was calculated using the motor age of onset and CAG repeat length. To subtract the effects of CAG repeats from the age of onset, a previously developed phenotype model was utilized. To mitigate skewness in the distribution of residual age at onset and to model a theoretical normal distribution, a Box-Cox power transformation was applied to the distribution prior to its usage as a phenotype for association testing. The resulting significance values from the GWA were uploaded to the LocusZoom browser and plotted for the rs3812963 locus.

### Patient-derived neuronal cultures

Striatal medium spiny neurons (MSNs) were directly converted from the fibroblasts of healthy controls and HD patients using brain-enriched microRNAs, miR-9/9* and miR-124 and subtype-defining transcription factors CTIP2, DLX1/2, and MYT1L as previously described^74, 108^. Three independent cell lines were used. P2A astrocytes and MT3 astrocytes were co-cultured with Control MSNs (Ctrl-MSNs - n=1 cell line) and HD-MSNs (n=3 cell lines) at 1:1 ratio on post-induction day 22 (PID22). Reprogrammed cells were treated with Incucyte Caspase-3/7 Green Reagent and Annexin V Red Reagent on PID27. 4 to 6 technical replicates per condition were used. Imaging scheduling, collection and data analyses were performed with the Incucyte Live-Cell Analysis System. Ctrl-MSNs and HD-MSNs were imaged every 24hours for 8 days (PID28 to 35). Images were analyzed for the number of green or red objects per well of 96 well plates. For the apoptotic index, the number of green or red objects divided by phase area (µm2) per well was quantified.

### Co-Cultures of astrocyte with murine neurons

Murine Neuro 2a (N2A - Sigma cat # 89121404) were cultured in DMEM + 10% FBS according to vendors protocols. The cells were seeded onto a cell culture treated 24 well-plate at 3.5X10^4 cells/well in for 24hours. The next day, the media was changed and switched to a DMEM differentiation media containing 2% FBS and 20 uM of Retinoic acid. Cells were allowed to differentiate for 24 hours before adding control GFP or MT3 overexpressing astrocytes at 3.5X10^4 cells/well in astrocyte culture medium (Sciencell©). The cells were co-cultured for 24 hours before adding Cadmium at 50uM, 10uM or Rotenone at 200nM or 20nM, versus DMSO. 24 hours later, the cells were stained with Propidium Iodide (Invitrogen cat# P3566) at 1:500 for 30 minutes prior to washing, trypsinization, and analysis by flowcytometry using BD Bioscience LSRII flowcytometer. After gating on the live singlet cells, the percentage of positive FITC+/- that were PI+/- were quantified by FCS express 7 (De Novo Software). The experiment was replicated 4 times.

### Astrocyte-microglia co-culture for microglia RNAseq and phagocytosis assay

HMC3 cells (ATCC cat#CRL-3304) were seeded in 24 well-plate at 5X10^4 each well in 0.5ml culture medium (DMEM with 10% FBS medium) and cultured overnight before co-culture with control, CLU overexpressing, or MT3 overexpressing astrocytes in a transwell assay. The astrocytes were seeded in the upper chamber of a 24 well-plate 0.4um polycarbonate membrane inserts (costar cat#3413) pre-coated with Poly L-Lysine, at 5X10^4 cells per insert in 0.1ml of astrocyte growth medium (Sciencell™) 3 hours before transferring the insert to the wells containing the microglia. The two cell types were co-cultured for 24hrs before the astrocyte inserts were removed for collecting microglial RNA for RNAseq or conducting the phagocytosis assay. For the latter, the media was changed into fresh HMC3 media containing fluorescent latex beads (1:500, latex -eads-rabbit IgG-FITC complex - Cayman chemical cat#500290). The cells were incubated for 30 min at 37°C before being washed as the manufacturer protocol, trypsinized, and analyzed by flowcytometry using BD Bioscience LSRII flowcytometer. The percentage of positive FITC+ cells was evaluated by FCS express 7 (De Novo Software) after gating on the live singlet cells. The experiment was replicated 4 times.

## Data availability

The human snRNAseq and bulk RNAseq dataset will be available in GEO under accession GSE242198 (reviewer tokens are available here: https://nam02.safelinks.protection.outlook.com/?url=https%3A%2F%2Fwww.ncbi.nlm.nih.gov%2Fgeo%2Fquery%2Facc.cgi%3Facc%3DGSE242198&data=05%7C01%7Cfp2409%40cumc.columbia.edu%7C17522f482d974df9376108dbab24ce07%7Cb0002a9b0017404d97dc3d3bab09be81%7C0%7C0%7C638291946793497114%7CUnknown%7CTWFpbGZsb3d8eyJWIjoiMC4wLjAwMDAiLCJQIjoiV2luMzIiLCJBTiI6Ik1haWwiLCJXVCI6Mn0%3D%7C3000%7C%7C%7C&sdata=jHWHt08tlawBSgAWjERFlKqG4DgDSbnNzJQfLnxPSxA%3D&reserved=0). All other datasets are provided in the supplementary material and/or available from the corresponding author upon request. The customized code used for analysis in this manuscript is provided here: https://github.com/Al-Dalahmah-lab/HD_astrocytes_paper.

## Author contribution

FP, AY, JEG, VM, and OA designed the study. OA, AT, KO, NM, HL, JL, JSK, AM performed the experiments. JSK and AY perform the HD patient neuronal cultures experiments. CWN, AB, DEH, CH conducted the GWAS analysis. HL performed cell culture and flowcytometry analysis. JL, NM, AT performed the sequencing experiments. KO performed the WGCNA analysis. OA, FP, NM, VM, JSK, AT, JEG, CN analyzed the data. All authors read and approved the manuscript.

## Acknowledgements

OA was supported previously through funding from the Huntington Disease Society of America and the Hereditary Disease Foundation, and Currently by the American brain tumor association, NIA ADRC REC program Grant Number P30AG066462. OA, FP, JEG and VM are supported by NIH R21 AG075754. This research was supported by the Digital Computational Pathology Laboratory in the Department of Pathology and Cell Biology at Columbia University Irving Medical Cancer, and by the Biomarkers Core Laboratory at the Irving Institute for Clinical and Translational Research, home to Columbia University’s Clinical and Translational Science Award. We thank Nancy Wexler for all her support throughout this project.

## Supplemental results

### Weighted gene correlation network analysis reveals loss of preservation and connectivity of glial and immune modules

We clustered the bulk RNAseq samples as a preliminary analysis to uncover the main factors that drive gene variation in our dataset. Clustering of the samples based on a distance metric calculated from normalized gene expression identified 6 clusters, some with predominance of cortical samples (Cl3 and Cl5), striatal samples (Cl2, Cl4, Cl6), control samples (Cl3-4), or HD samples (Cl2 and Cl6 - **Figure S1B**). Clusters Cl2 and Cl4 were enriched in samples from females and males, respectively. These results show a strong influence of brain region and condition on gene expression, as expected.

To understand patterns of gene co-regulation and dysregulation in HD brain regions, we performed weighted gene correlation network analysis (WGCNA) on control and HD bulk RNAseq samples. Networks from control and HD samples were constructed separately to examine the changes in the patterns of gene correlation in HD (**Figure S2A**) as described in the *Methods*. Gene comprising each module are provided in **Supplementary Table 3**. 22 and 20 modules were identified in the control and HD networks, respectively (**Figure S2A**). Multiple control modules exhibited gene overlap with HD modules, however, some HD gene modules appeared to show minimal overlap with control modules - for instance, the HD purple and turquoise modules (**Figure S2B**), suggesting that genes in HD gain correlation. We next correlated the genes in the control and HD modules to anatomic region, sex, age, and CAG repeat length (in HD modules) and found that there were several control modules with significant, strong, positive and negative correlations with anatomic regions (**Figure S2C**), and CAG repeat length (in HD modules). In particular, the HD purple module (which did not exhibit significant overlap with control modules) and the lightgreen module were significantly positively correlated with CAG repeat length, and the black, blue, turquoise, and lightcyan were significantly negatively correlated with CAG repeat lengths. Next, we asked which control modules lose preservation in HD. Examination of preservation statistics reveals that several modules showed minimal preservation overall, and in particular, loss of density and connectivity (**Figure S2D** **and Supplementary Table-3).** The loss of connectivity of select modules is shown in **Figure S2G**. These modules include the lightgreen, darkred, pink, tan, lightyellow, and royalblue. The former (lightgreen) is positively correlated with the caudate region, while the others are negatively correlated with the caudate and display cingulate (darkred) or accumbens (tan) correlations. The royalblue and turquoise showed no significant trait correlations, which suggests they correspond to regional-agnostic gene programs. Examination of gene ontologies enriched in the most poorly preserved modules incriminate splicing, meiotic DNA double-strand break processing, DNA repair, mitosis, transcription, nuclear export, T-helper cell lineage commitment, antigen presentation, carbohydrate metabolism, response to iron ions, dopamine metabolism, and synaptic function (**Figure S2E** and **Supplementary Table-3**). Next, we examined the GO terms that are enriched in the HD modules that were significantly positively correlated with CAG repeat length - including the lightgreen and purple. The results incriminate enrichment of genes involved in immune response, T cell function, translation, proteosomal function, polyubiquitination, cilium function, development, response to stress, and response to DNA damage. Of the genes enriched in the modules negatively correlated with CAG repeat length, our results showed enrichment of genes involved with vascular function and angiogenesis (lightcyan), base-excision repair, histone methylation, and T-cell receptor signaling (blue), and response to unfolded protein, RNA splicing, cell aging, and myeloid cell homeostasis (black - **Figure S2F** and **Supplementary Table-**3). Given the involvement of immune genes in modules poorly preserved in HD, we asked if modules associated with immune (microglial) and astroglial module genes display altered connectivity. Similar to poorly preserved modules (lightgreen, lightcyan, darkred, tan, pink), hub genes in the control black (immune) and brown (astroglial/immune) modules showed loss of connectivity of the hub genes, which appear to be members of other modules in the HD network (**Figure S2G**). Together, these results confirm that HD indeed significantly influences astrocyte and microglial gene expression.

### A subset of fibrous-like astrocytes expresses DPP10

One of the genes abundantly expressed in our fibrous-like astrocyte dataset is DPP10. This encodes a dipeptidyl peptidase enzyme and voltage gated potassium channel increased in neurodegenerative diseases^1^. This gene is expressed mainly in neurons and we asked if it is expressed in astrocytes. Thus, we performed dual CD44 DPP10 labeling of the caudate nucleus of 3 controls and 3 HD brains (**Figure S4C**). In all cases, we found the protein to be co-expressed with CD44 in a small subset of CD44 positive astrocytes in the parenchyma (HD), pencil fibers, subependymal zone, and around large blood vessels.

### Differentially expressed genes in neurons

We performed differential gene expression analysis on major neuronal cell types detected in the striatal and cortical regions of our study. Volcano plots displaying top DEGs for each cell type in all three brain regions are displayed in **Figure S7A-C**. The differential gene expression analysis of neurons revealed that the overall numbers of DEG was significantly larger in the striatal region than the cortex (**Figure S7D**). We saw some of the highest numbers of DEG in the SPN cells with the accumbens dSPN having the largest number of DEG decreased and the highest upregulation coming from the caudate dSPN. In the cingulate cortex, the glutamatergic neurons had significantly more DEG compared to the GABAergic neurons and the number of DEGs from the GABAergic neurons were significantly lower compared to the striatal regions. The DEGs are provided in **Supplementary Table 8**.

To delve into the details (**Supplementary Table-8**), we found the expected decrements of PCP4 and PDE10A in all SPNs in the caudate and accumbens. In dSPNs of both regions, we found the expression of PENK to be increased. Changes in these genes have been documented by multiple studies ^2–4^, to name a few. Moreover, the caudate dSPN showed a significant decrease in the expression of TAC1 but this was not found to be significant in the accumbens. Among the set of genes that were shared and downregulated between SPNs across the two regions were MALAT1, FTX, RBFOX1, RYR3, and RYR2. Interestingly, we observed that the expression of CLU to be increased in dSPN and iSPN neurons in the accumbens and caudate along with a significant increase in accumbens GABAergic cells as well.

We compared our DEG results from the caudate to Lee et al.^5^ SPN DEG. We found 291 genes in common among dSPN in the caudate that were upregulated in our analysis and 251 genes that were downregulated (**Supplementary Table 8**). Many of the genes that were upregulated were involved in heat shock protein response and immune response. More specifically, these genes included HSB1, HSPA4, HSPA8, HSPA1A, CD81, and CLU. Genes that were downregulated in dSPN included SLC24A2, RYR3, CACNA1A, JPH4, and CALM1 which are known to be involved in calcium ion activity. Comparing the iSPN neurons to the Lee et al. findings revealed there were 210 shared genes that were upregulated and 327 shared genes that were downregulated. Many of the shared genes that were upregulated were similar to the dSPN like HSPA1A, CLU, DNAJA1. Among the upregulated genes, a large portion of it were involved in responding to unfolded proteins. In addition, we examined the medium spiny neuron DEG from Malaiya et al^4^ and confirmed that many of the genes reported to be downregulated across all MSN subtypes were indeed downregulated in dSPN and iSPN in the striatal brain regions of our data too. These genes include PDE10A, PDE1B, ADCY5, ATP2B1, and ARRP21. Shared upregulated genes in the iSPN are largely involved in response to unfolded protein which are driven by heat shock protein genes. On the other hand, some shared downregulated genes in the iSPN were involved with disruption of ion channel activity as driven by the decreased expression of genes like GRIN2A, GRIA1, GRIN2B, NLGN1, DAPK1, DLG1, and DLG2.

We also compared our GO terms to the Lee et al. and found that our results are consistent with the decrease of phosphatidylinositol signaling system in accumbens dSPN and iSPN as well as in the caudate dSPN but not caudate iSPN. MAPK signaling was only found to be increased among accumbens dSPN. Conversely, the GO Cholinergic synapse was increased in Lee et al. but decreased in all of our SPN in accumbens and caudate. Spliceosome activity also was decreased as expected among all SPN in all regions. These differences may arise from differences in the regions analyzed in Lee et al, which included the putamen - a region not included in our analysis. In addition to these comparison, we also report genes involved in various ion channel activity as well as lipid phosphorylation to be depleted in all SPN in both the accumbens and caudate (**Figure S7E**). There were significant differences between the caudate and accumbens SPNs, and between iSPNs and dSPNs. For example, dendrite morphogenesis GO was increased in accumbens SPN but decreased in the caudate. Moreover, glutamatergic synapse GO term was significantly enriched among dSPN in both the accumbens and caudate but not in the iSPN. Also, GABA neurons showed fewer DEGs than other neuronal types, and even these were regionally diverse, being mildest in the cingulate cortex (**Figure S7D**). Among the cingulate neurons, we observe a general decrease in genes related to metabolism among the glutamatergic and GABAergic neurons.

### Astrocytes-Microglia cross-talk in HD

Given the known cross-talk between astrocytes and microglia^6^, we asked if microglial gene expression is altered in HD brains. Notwithstanding the limitations of snRNAseq in analyzing microglial cells^7^, and the relative paucity of retrieved cells, we went ahead and analyzed HD and control myeloid cells in our cohort. We first subclustered our myeloid cells from the three brain regions analyzed. We discovered 3 major clusters which correspond to microglial cells, blood derived myeloid cells (monocyte-derived macrophages), and T-cells (**Figure S10A**). This is further illustrated by examining gene marker expression (**Figure S10B** **and Supplementary Table-9)**. Since the number of macrophages and T-cells were comparably lower in numbers than microglial cells (**Supplementary Table-9**), we decided to focus on the latter for downstream analysis. Microglial cells were distributed across all brain regions examined, unlike T-cells and macrophages, which we mainly retrieved from HD striatal samples (**Supplementary Table-9**). To better understand how the microglia differ across the three brain regions in HD, we performed DEG analysis in each of the regions separately (**Figure S10C-E**). Interestingly, heat shock protein gene expression varied between the three brain regions in microglia. For example, HSPH1 was increased in the accumbens and cingulate but downregulated in the caudate. In addition to HSPH1, there were several other genes that were among the most downregulated in the caudate like HSPA1A, HSPA1B, HSP90AA1, and HSPH1. Examining the GO enriched in microglial DEGs showed regional differences in microglial responses to HD (**Figure S10F**). Cingulate and accumbens HD microglia upregulated genes involved in heat shock and stress response, but this was not the case in the caudate. Likewise, in the same regions, HD microglia upregulated genes associated with cell scavenger receptors, which are associated with the innate immune response and phagocytosis^8^.

Given the heterogeneity of gene expression changes in microglia in HD, and in light of our previous results on astrocyte phenotypic heterogeneity across brain regions in HD, we asked if specific astrocytic phenotypes that were region-enriched altered microglial function. Specifically, we examined if the neuroprotective astrocytic phenotype, with increased MT3 expression, led to changes in microglial phagocytosis. To address that question, we performed co-culture assays between microglia and astrocytes in a trans-well assay, and quantified microglial phagocytosis of fluorescent beads using FACS or gene expression in microglia using RNAseq (**Figure S11A**). We overexpressed MT3 and CLU (which were both markers of neuroprotective protoplasmic clusters 1 and 3), or GFP in astrocytes, and co-cultured with control microglia. We were interested in CLU because we had published a paper showing that CLU was increased in astrocytes in the context of glioma microenvironment, and that CLU+ astrocytes can alter gene expression of glioma cells in a manner that reduced expression of immune-related genes^9^. This experimental design allowed us to measure how changes in astrocyte phenotype influence microglial function via factors secreted into the media by astrocytes. Our results showed that co-culture of astrocytes with microglia significantly altered gene expression in microglia (**Figure S11B** - see PCA, and **Supplementary Table-9)**. However, while MT3 overexpression significantly altered gene expression in microglia (**Figure S11D**), that was not the case in microglia co-cultured with CLU overexpressing astrocytes (**Figure S11C**). MT3-astrocytes caused microglia to upregulate genes associated with lipid and fatty acid metabolism and transport (**Figure S11E**). Functionally, microglia co-cultured with MT3 but not CLU overexpressing astrocytes increased their phagocytic activity (**Figure S11G**). This suggests that neuroprotective astrocytes which express high levels of MT genes can increase phagocytosis in microglia, which we posit is a compensatory “positive” phenomenon. Finally, driven by these changes in microglial gene expression induced by MT3 astrocytes, we asked if the genes significantly increased in microglia as induced by MT3 astrocytes (referred as MT3-astro signature) were enriched in the DEGs in HD vs control microglia from our snRNAseq data across the three brain regions. We found that while the MT3-astro signature was significantly enriched in the cingulate and approached significance in the nucleus accumbens, it was not enriched in the caudate (**Figure S10F**). This was a welcome finding given that MTs were significantly increased in the cingulate and accumbens protoplasmic astrocytes (by snRNAseq and IHC validation for cingulate, snRNAseq for accumbens), but not in caudate astrocytes (by IHC analysis). Together, these findings suggest that in addition to the neuroprotective and Astro-protective roles of MTs, these proteins also lead to significant changes in microglial gene expression and increased phagocytosis, which are most likely compensatory and favorable.

## Supplementary figure legend

**Figure S1.**
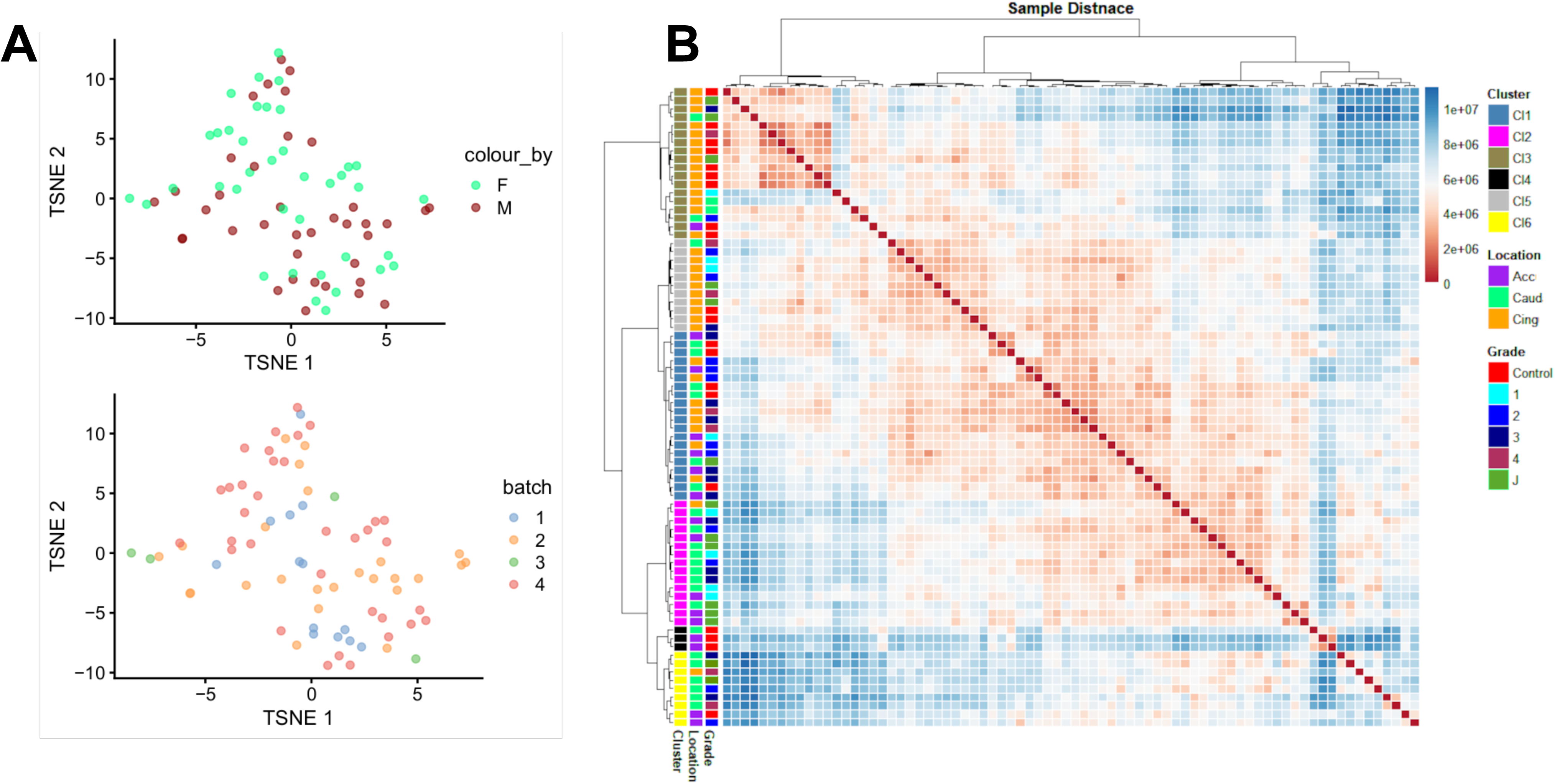
Related to Figure 1. **A**) t-distributed stochastic neighbor (tSNE) embedding of bulk RNAseq samples used in the study color-coded by sex (top), and sequencing batch (bottom) - showing adequate correction for batch- and sex- related effects. **B**) Sample distance heatmap showing the Manhattan distance clustered using the Ward D2 method. Normalized batch corrected counts were used to generate the sample distance. The colored bars on the left indicate the sample clusters, anatomic region (Acc: Accumbens. Caud: Caudate, Cing: Cingulate), and HD grade/Condition (Control (Con) versus HD grades 1-4 & J: juvenile onset HD).

**Figure S2.**
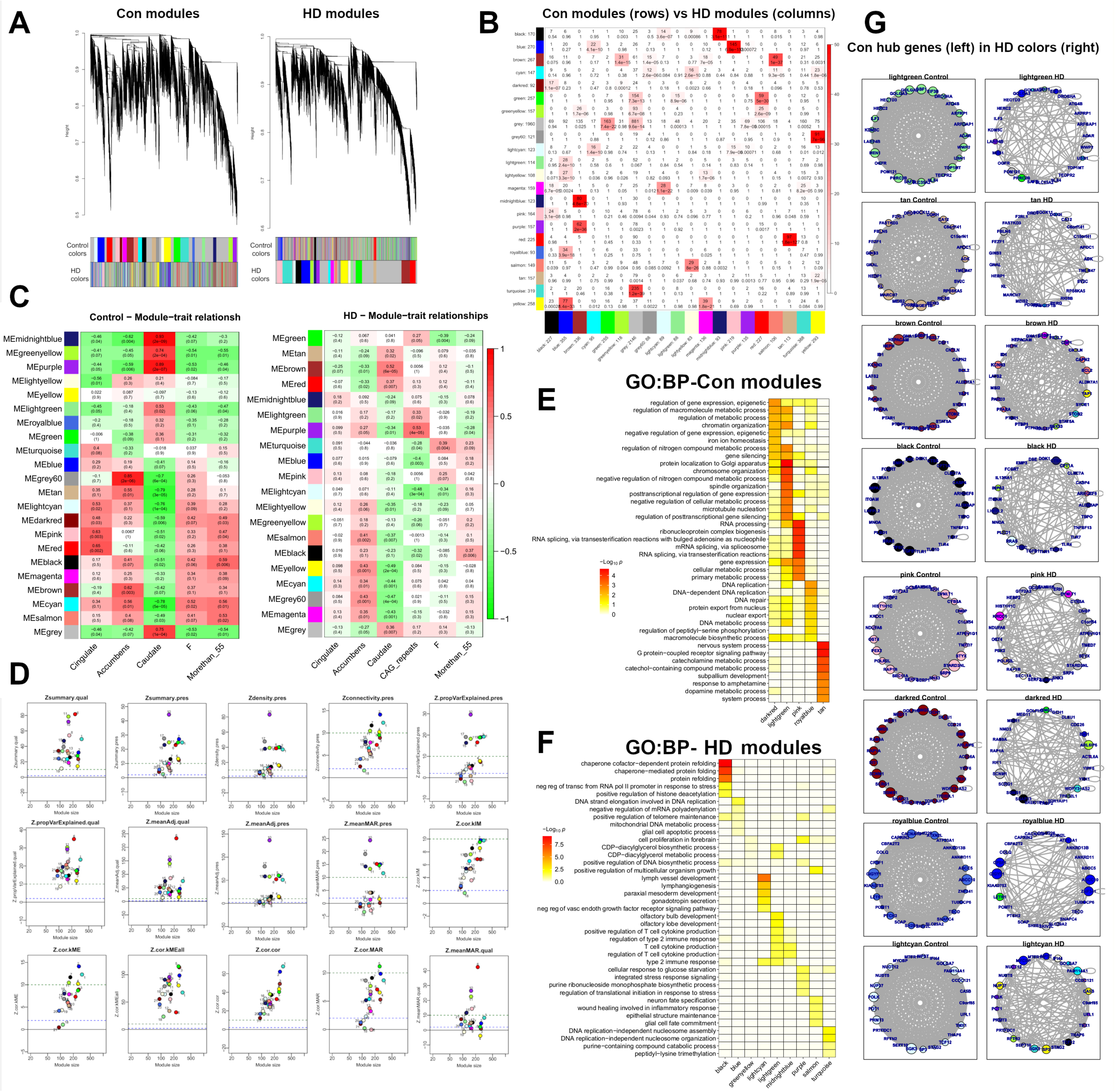
Weighted gene correlation analysis (WGCNA) identifies gene modules that loose preservation in HD. **A)** Dendrograms showing genes clustered hierarchically for control **(right)** and HD **(left)** samples. The genes are colored by control as well as HD network module designations, and vice versa, in the respective control (left) and HD (right) dendrograms. **B**) Heatmap showing the overlap between genes in control modules (rows) and HD modules (columns). The number of overlapping genes and p values are indicated in the tiles. **C**) Heatmap of trait-module correlations for the Control (left) and HD (right) network modules displaying correlation coefficients (and p values in parenthesis). The module-trait relationships were determined in control and HD expression matrices, separately. **D**) Control module preservation statistics plotting module size on the x-axis against several module density and connectivity statistics on the y-axis. The top row shows the most frequently used summary statistics. The color of the nodes represents gene module membership in control and HD modules. **E**-**F**) Gene Ontology (GO) term enrichment analysis showing the top biological process GO terms enriched in the genes of select control (**E**) and HD (**F**) modules. The heatmap represents the GOs as rows, and module names as columns. The negative log10 of the enrichment p value is represented by color. **G**) Subnetwork analysis of the top 25 genes with highest connectivity (hub genes) in the control modules shown in the control subnetwork (left) versus the HD subnetwork (right). Nodes represent genes and edges correlations. The edges with weights less than the mean weight for each subnetwork were trimmed. The size of the circles (gene) represents the hub score as detected in igraph package. The node color in the HD column (right) indicate the module membership of the respective gene in the HD network.

**Figure S3.**
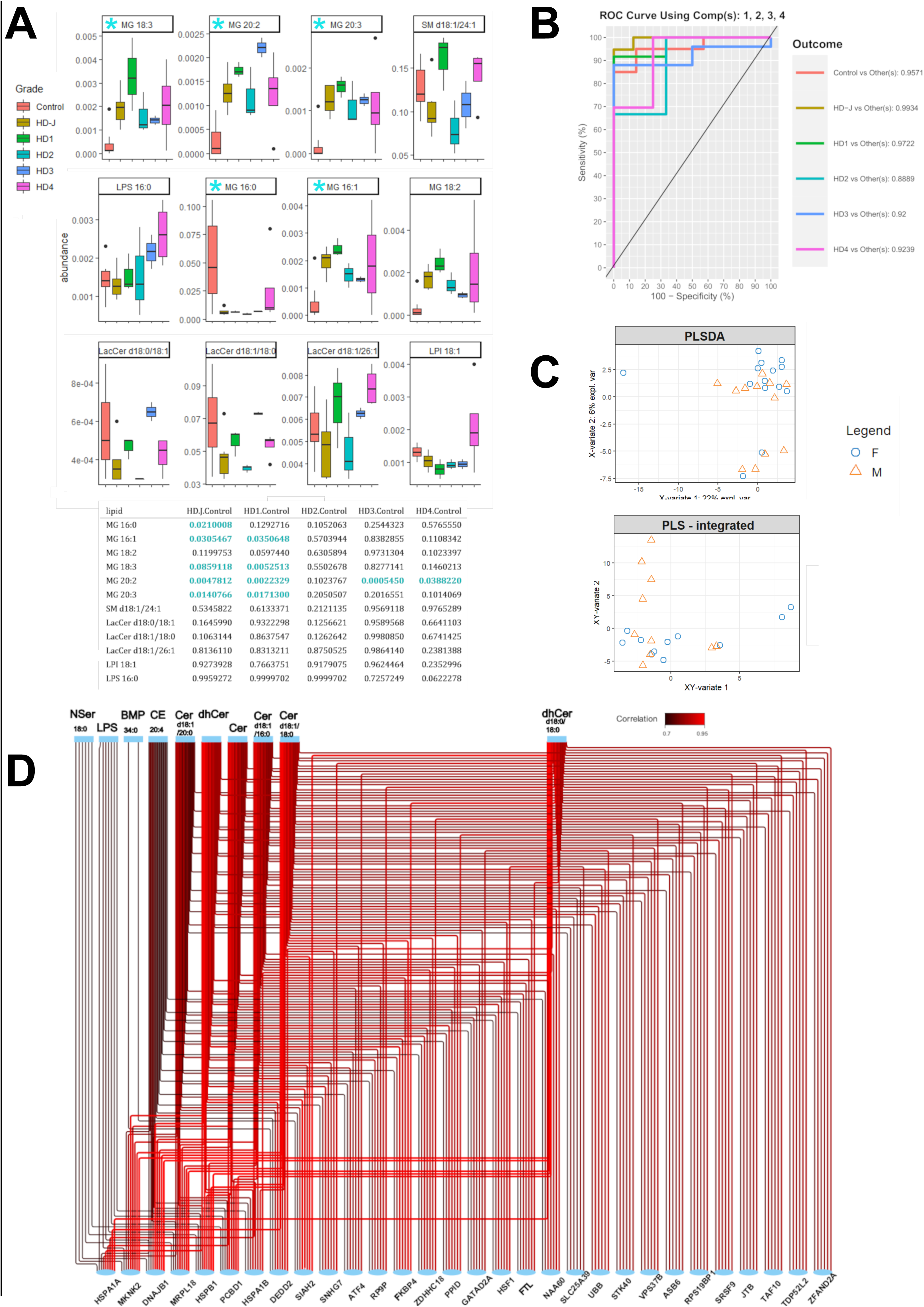
Related to Figure 2. **A**) Boxplots of relative abundance of select long-chain lipid species with significant one-way ANOVA p-values for grade. The data is presented with grade (color-coded) on the x-axis. The p-values for Tuckey post-hoc comparisons are indicated in the table (down). Stars indicate significant post-hoc analysis p-values. **B**) Receiver Operating Characteristic (ROC) curve showing the sensitivity (y-axis) and 100-specificity (x-axis) of the sPLS-DA model, based on 4 components, to distinguish condition (grade). ROC traces for each grade are represented separately and denoted using the colors indicated on the right. **C-D**) Scatter plots showing projection of lipidomics samples and integrated lipidomic/RNAseq in the first two latent variables of sPLS. The variance explained by each component is indicated on the axes. The samples are color- and shape- coded by sex. **D**) Correlation network analysis showing the correlation (encoded by edges, with color corresponding to correlation values) between genes and lipid species (nodes). Only the most highly (positively and significantly) correlated nodes are shown.

**Figure S4.**
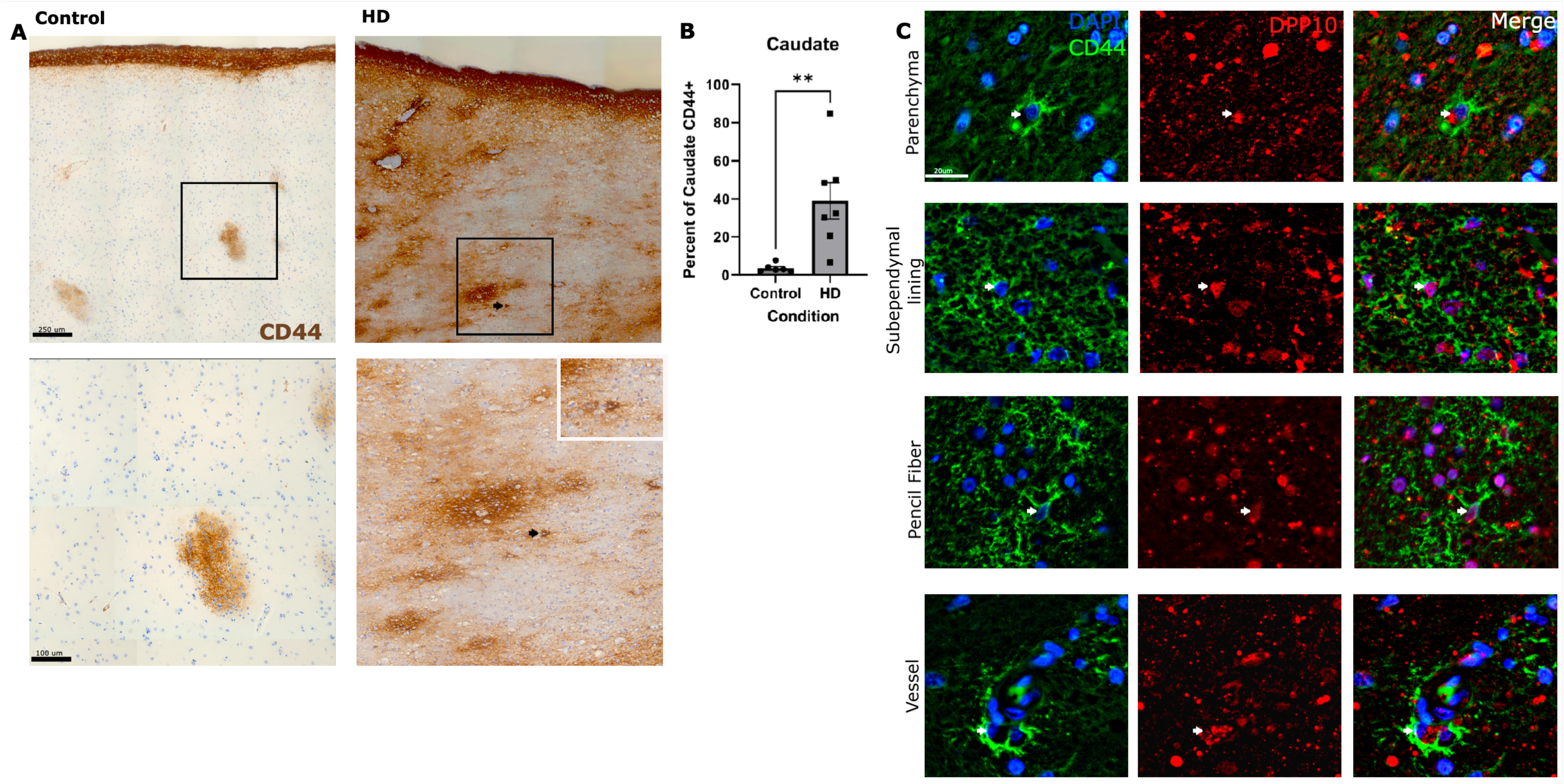
Increased CD44 expression in the Caudate of HD astrocytes. **A)** Immunohistochemical stain for CD44 in the Caudate of control and HD samples. Note the labeling of pencil fibers and ventricle lining seen at the top of the image. The arrow indicates a CD44 positive cell in the caudate parenchyma not associated with large vessels or pencil fibers. Scale bar= 250µm. Inset - enlarged image of the region shown in the black box in A showing the parenchymal CD44 positive cell in the top right corner of the HD image. Scale bar=100µm. **B)** Quantification of the percent of the ROI that was positive for CD44 using a one-tailed unpaired t-test. N=6 for the control and 7 for HD. Data is shown as mean +/- SEM. P value= 0.0029. **C**) Double immunofluorescence of CD44 (green) and DPP10 (red) in different caudate brain regions as indicated on the left. Scale bar=20µm.

**Figure S5.**
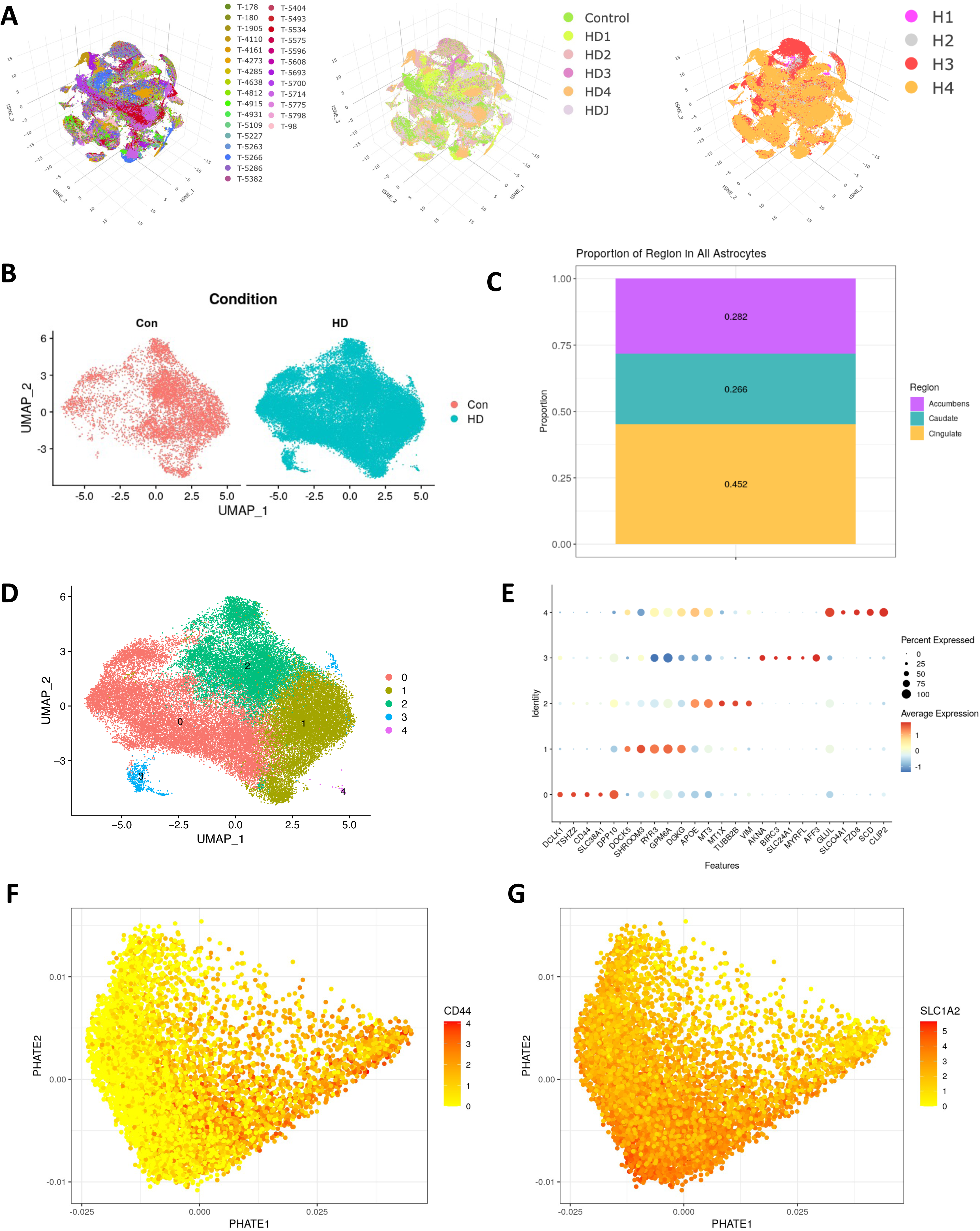
Related to Figure 3. **A**) t-SNE plots of all snRNAseq cells color-coded by donor (left), grade (middle), and sequencing batch (right). **B)** Split UMAP plots of astrocytes by condition where the left plot is control cells, and the right is the HD cells. **C)** Bar plot showing the proportion of each region among all the astrocytes. **D)** Clustering of all astrocytes as shown in the UMAP space. **E)** Dot plot showing the expression of the top five cluster markers for the clusters in (**D**). **F**) **Related to** **Figure 5D**. PHATE embedding of fibrous-caudate astrocyte with each cell color-coded by normalized *CD44* expression. **G**) Same PHATE embedding in (**F**) but with *SLC1A2* expression.

**Figure S6:**
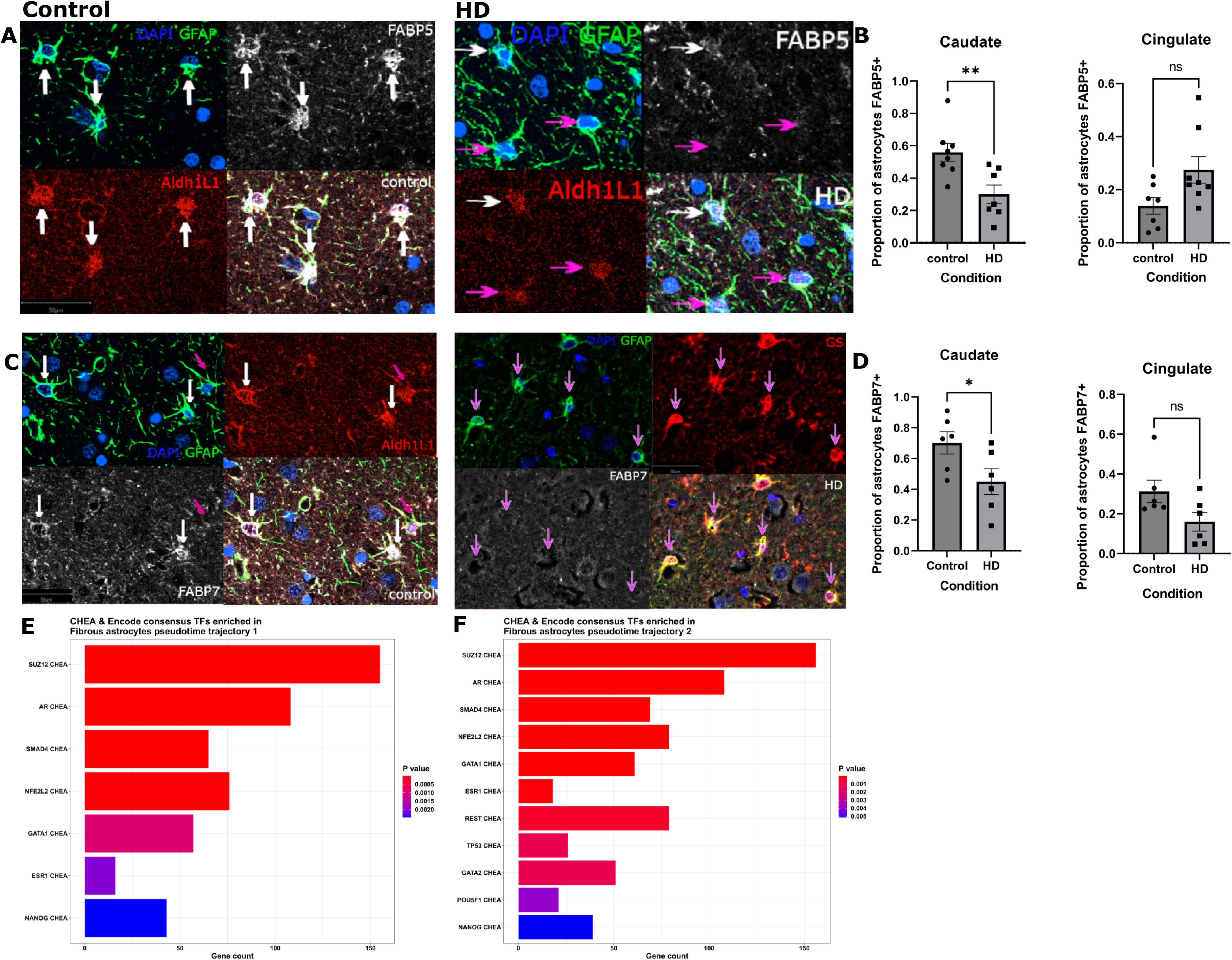
FABP5 and FABP7 expression are decreased in the caudate HD astrocytes. **A)** Immunofluorescent images of the caudate labeled for nuclei (DAPI-blue) and GFAP (green) to detect astrocytes (upper left), FAPB5 (white-upper right) and ALDH1L1 (red-bottom left). A merge of the four channels is shown in the bottom right panel. White arrows indicate DAPI, GFAP and FABP5 positive cells (FABP5 positive astrocytes) and pink arrows indicate FABP5 negative astrocytes. Scale bar=50µm. **B)** Quantification of the proportion of FABP5 positive astrocytes in the caudate. Two-tailed Mann-Whitney test used with N=8 for control and 7 for HD. Data is shown as mean +/- SEM. P value= 0.0093. **C)** Immunofluorescent images of the caudate labeled for nuclei (DAPI-blue) and GFAP (green) to detect astrocytes (upper left), FAPB7 (white-upper right) and ALDH1L1 (red-bottom left). A merge of the four channels is shown in the bottom right panel. White arrows indicate DAPI, GFAP and FABP7 positive cells (FABP7 positive astrocytes) and pink arrows indicate FABP7 negative astrocytes. Scale bar=50µm. **D)** Quantification of the proportion of FABP7 positive astrocytes in the caudate. Two-tailed Mann-Whitney test used with N=6 for control and HD. Data is shown as mean +/- SEM. P value= 0.0411. p values were not significant in the cingulate. **E-F**) EnrichR bar plots showing the enrichment of the transcription factor binding sites in genes that vary along fibrous-like astrocyte pseudotime trajectories 1 and 2 (note that we consider trajectory 2 to be associated with CD44 state transition). This analysis is based on the consensus ChIPseq ChEA dataset. The adjusted p value is indicated by color, gene count on the x-axis, and transcription factor on the y axis.

**Figure S7.**
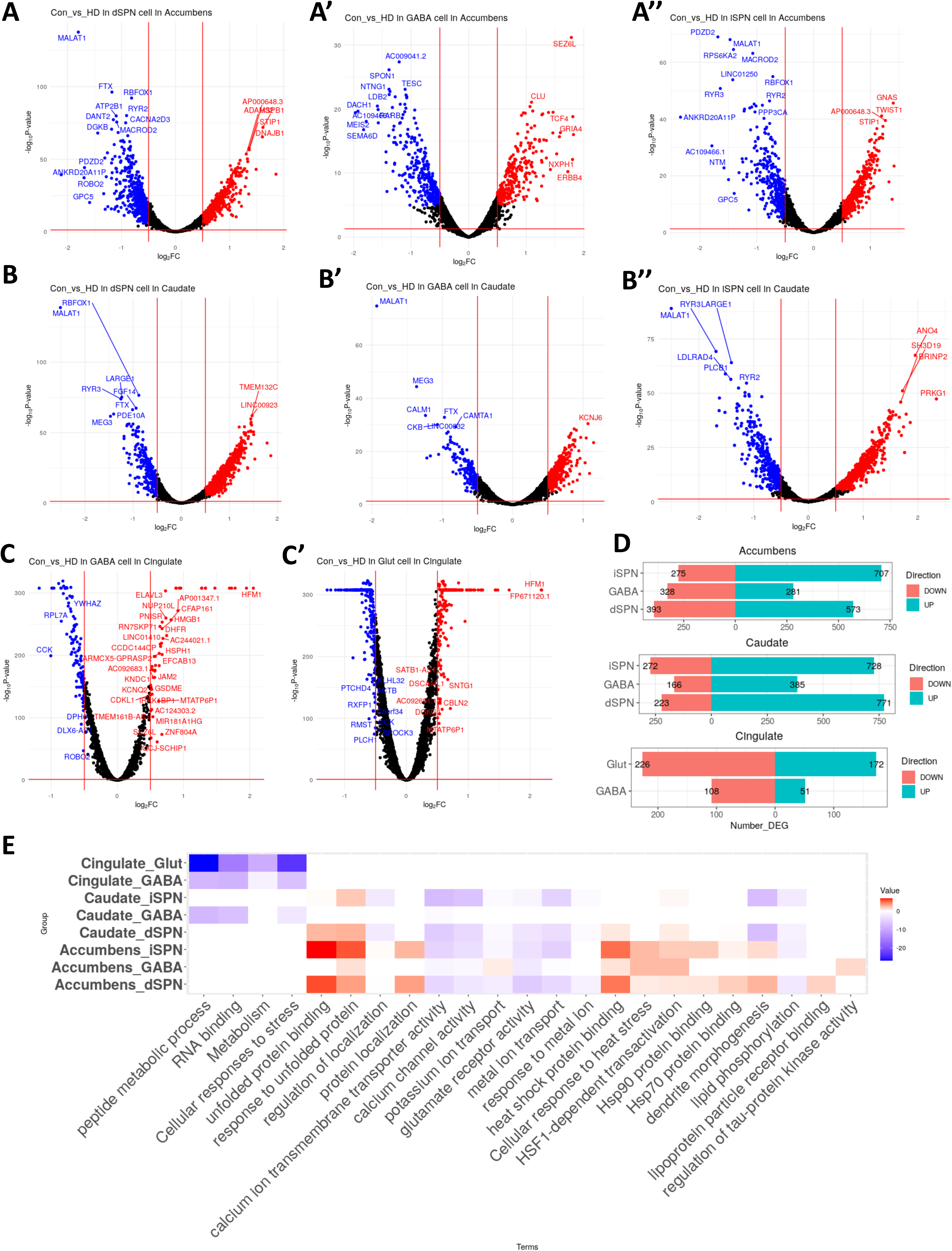
Differential gene expression in HD neurons. **A)** Volcano plots of genes from the accumbens neurons that are differentially expressed in HD (red highlight = genes significantly found to be higher in HD samples, blue highlight = genes significantly found to be lower in HD samples compared to control) within dSPN (direct-pathway spiny projection neurons), GABAergic neurons (**A’**), and iSPN (indirect-pathway spiny projection neurons) (**A’’**). **B)** Same analysis as (**A**) but for caudate neurons. **C)** Volcano plots for differentially expressed genes from the cingulate GABAergic neurons and Glut (Glutamatergic) neurons (**C’**). **D)** Bar plot displaying the number of DEG (based on the top 1000 genes, |logFC| > 0.5, and adjusted p-value < 0.05) for each major neuronal type across all three brain regions. **E)** Heatmap displaying the log10(p-value) of the GO terms (rows) enriched in DEGs comparing HD to control clusters (columns from **A-C**); red color indicates terms significantly enriched in DEGs increased in HD, and blue indicates terms significantly enriched in DEGs decreased in HD, and white means no significance.

**Figure S8.**
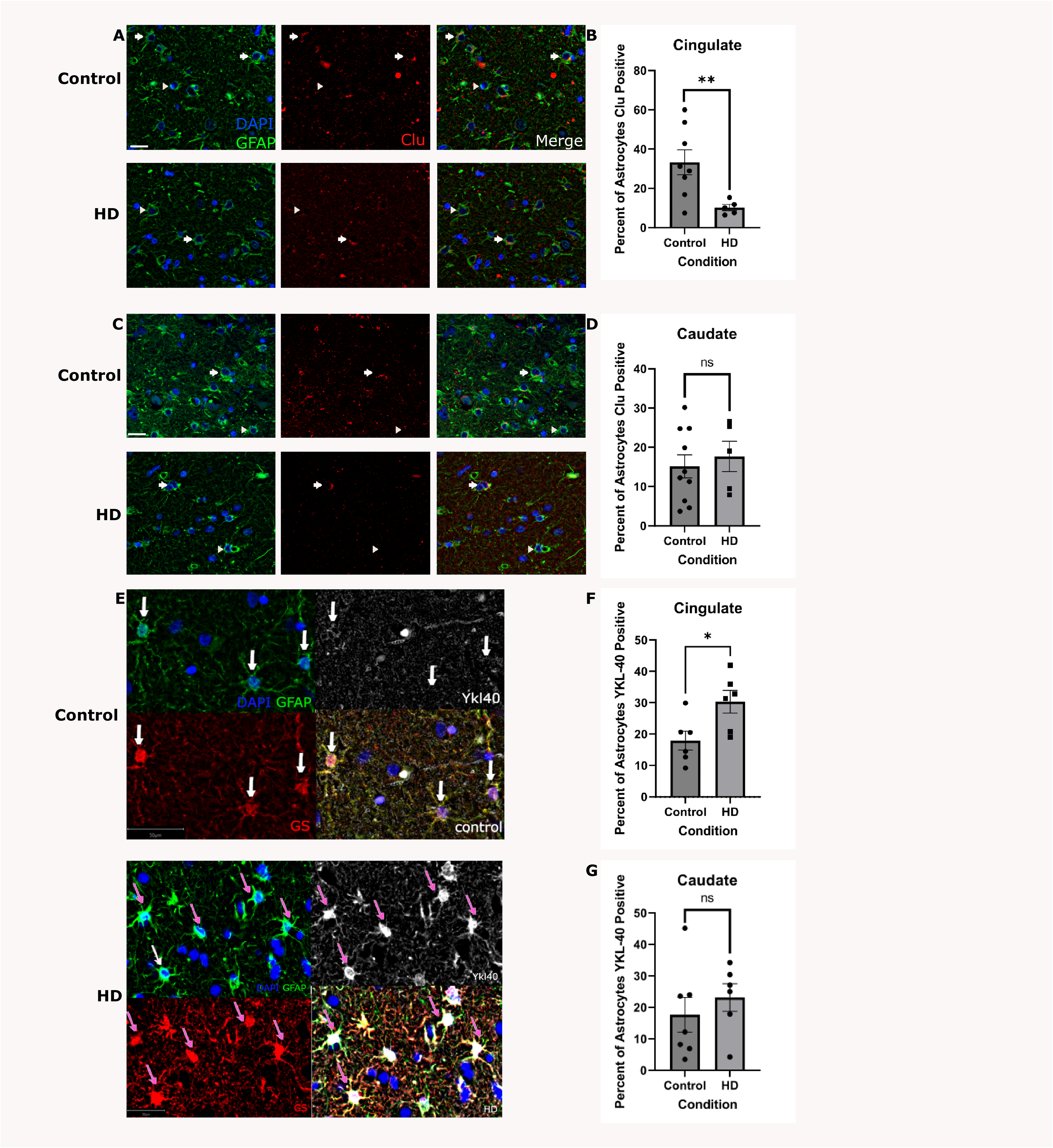
Decreased CLU Expression and Increased YKL-40 (CHI3L1) Expression in cingulate HD astrocytes. **A)** Representative multiplex immunofluorescence images of the Cingulate cortex labeled for nuclei (DAPI-blue) and GFAP (green) to detect astrocytes (left), and CLU (red-middle). A merge of the three channels is shown on the right. Arrows indicate DAPI, GFAP and CLU positive cells (CLU positive astrocytes) and arrowheads indicate CLU negative astrocytes. Scale bar=20µm. **B)** Quantification of the percent of CLU positive astrocytes in the Cingulate. Unpaired one-tailed T-test with N=8 for control and 6 for HD. Data is shown as mean +/- SEM. P value= 0.0086. **C)** Same as A but for the caudate. **D)** Quantification of the percent of CLU positive astrocytes in the Caudate. Unpaired one-tailed T-test used with N=10 for control and 5 for HD. Data is shown as mean +/- SEM. P value= 0.3130. **E)** Immunofluorescent images of the Cingulate labeled for nuclei (DAPI-blue) and GFAP (green) to detect astrocytes (upper left), YKL-40 (white-upper right) and GS (red-bottom left). A merge of the four channels is shown on the bottom right. Pink arrows indicate DAPI, GFAP and YKL-40 positive astrocytes and white arrows indicate YKL-40 negative astrocytes. Scale bar= 50µm. F) Quantification of the percent of YKL-40 positive astrocytes in the Cingulate. Two-tailed Mann-Whiteney test used with N=6 for control and HD. Data is shown as mean +/- SEM. P value= 0.0260. G) Quantification of the percent of YKL-40 positive astrocytes in the Caudate. Two-tailed Mann-Whiteney test used with N=6 for control and HD. Data is shown as mean +/- SEM. P value= 0.2949.

**Figure S9.**
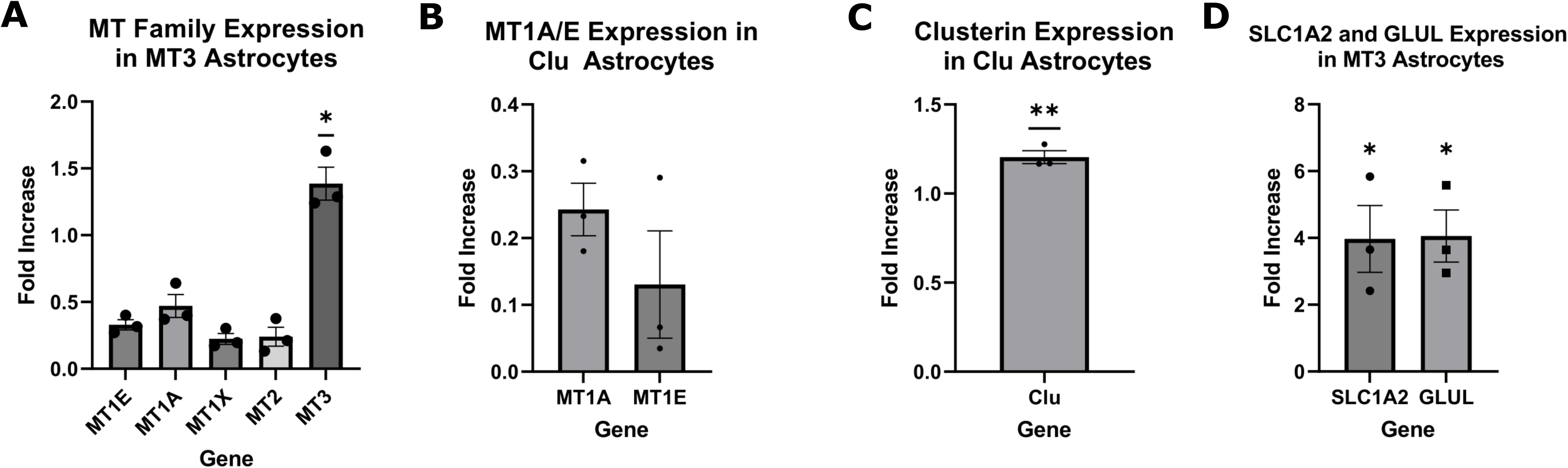
**A)** Gene expression quantification (real-time quantitative PCR rtPCR) of MT3 and other metallothioneins in MT3 overexpressing versus GFP control astrocytes. The delta-delta CT’s were log normalized. One sample T-test one-tailed test. P-value of 0.0125 for MT3 and non-significant for other metallothioneins. The gene expression was normalized for *GAPDH* control. **B-C)** Quantification of select MT family genes (**B**) and CLU (**C**) in CLU overexpressing astrocytes, The data is normalized and shown as per A. One sample two-tailed t-test, p-value= 0.1681 for *MT1E* and 0.2777 for *MT1A*, and p-value= 0.0016 for *CLU* in **C**. **D**) Gene expression quantification of *SLC1A2* and *GLUL* in MT3 overexpressing astrocytes by rtqPCR. The delta-delta CT compared to house-keeper (*GAPDH*) and control GFP astrocytes. One-tailed one-sample t-test. P values are indicated. MT3 overexpressing astrocytes significantly increased expression of *GLUL* and *SLC1A2.*

**Figure S10.**
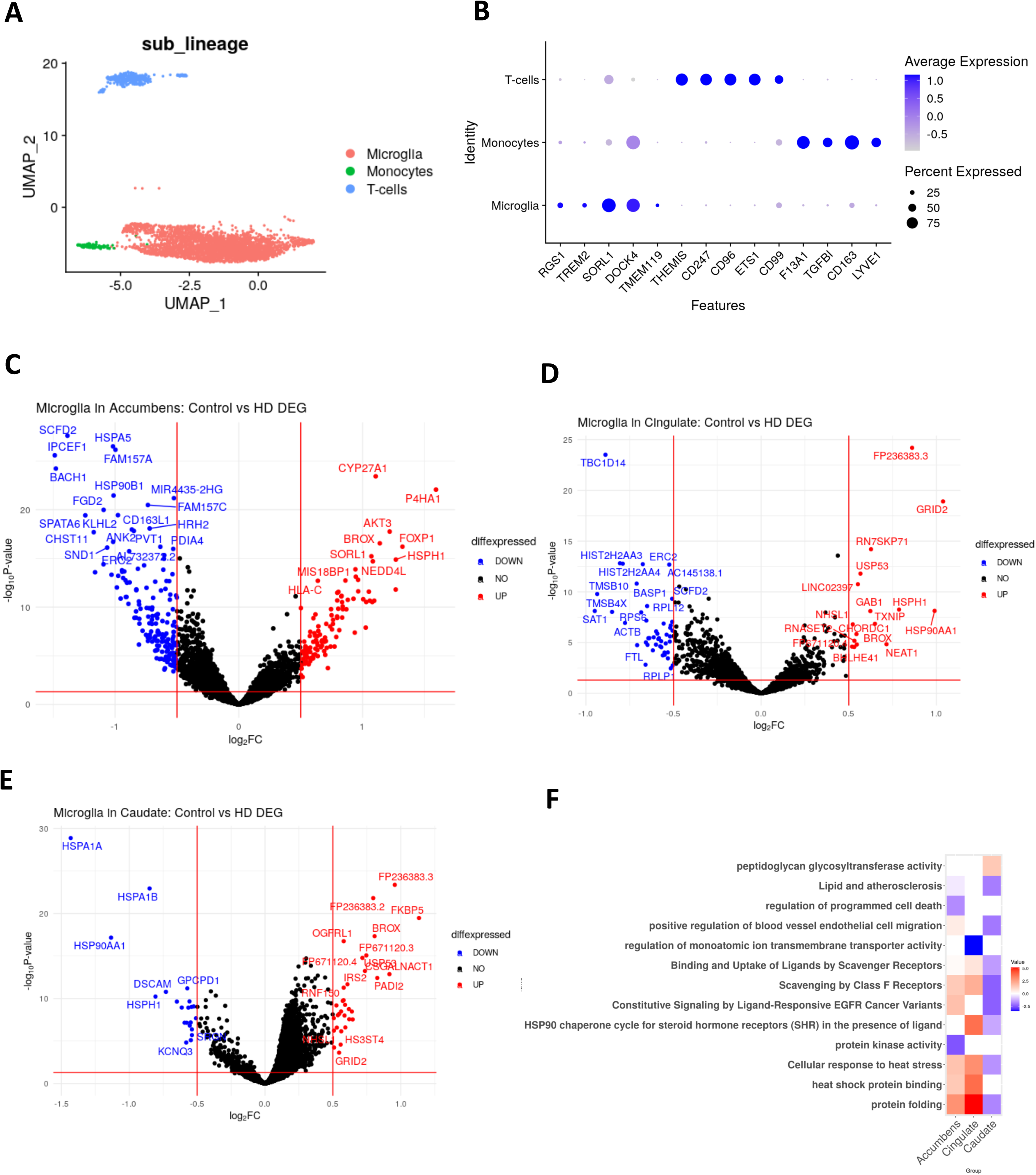
snRNAseq of myeloid population in HD and control brains. **A)** UMAP plot displaying the sub clusters of the main myeloid lineages. The three main cell types discovered are shown, microglia, T-cell, and monocytes. **B)** Dot plot showcasing markers for the three cell types found. **C)** Volcano plot of genes that are differentially expressed in HD for microglia found in accumbens (red highlight = genes significantly higher in HD samples, blue highlight = genes significantly lower in HD samples). **D)** Volcano plot similar to **(C)** but for microglia in the cingulate. **E)** Volcano plot similar to **(C)** but for microglia in the caudate. **F)** Heatmap displaying the log10(p-value) of the GO terms enriched in the DEGs from **(C), (D),** and **(E).**

**Figure S11.**
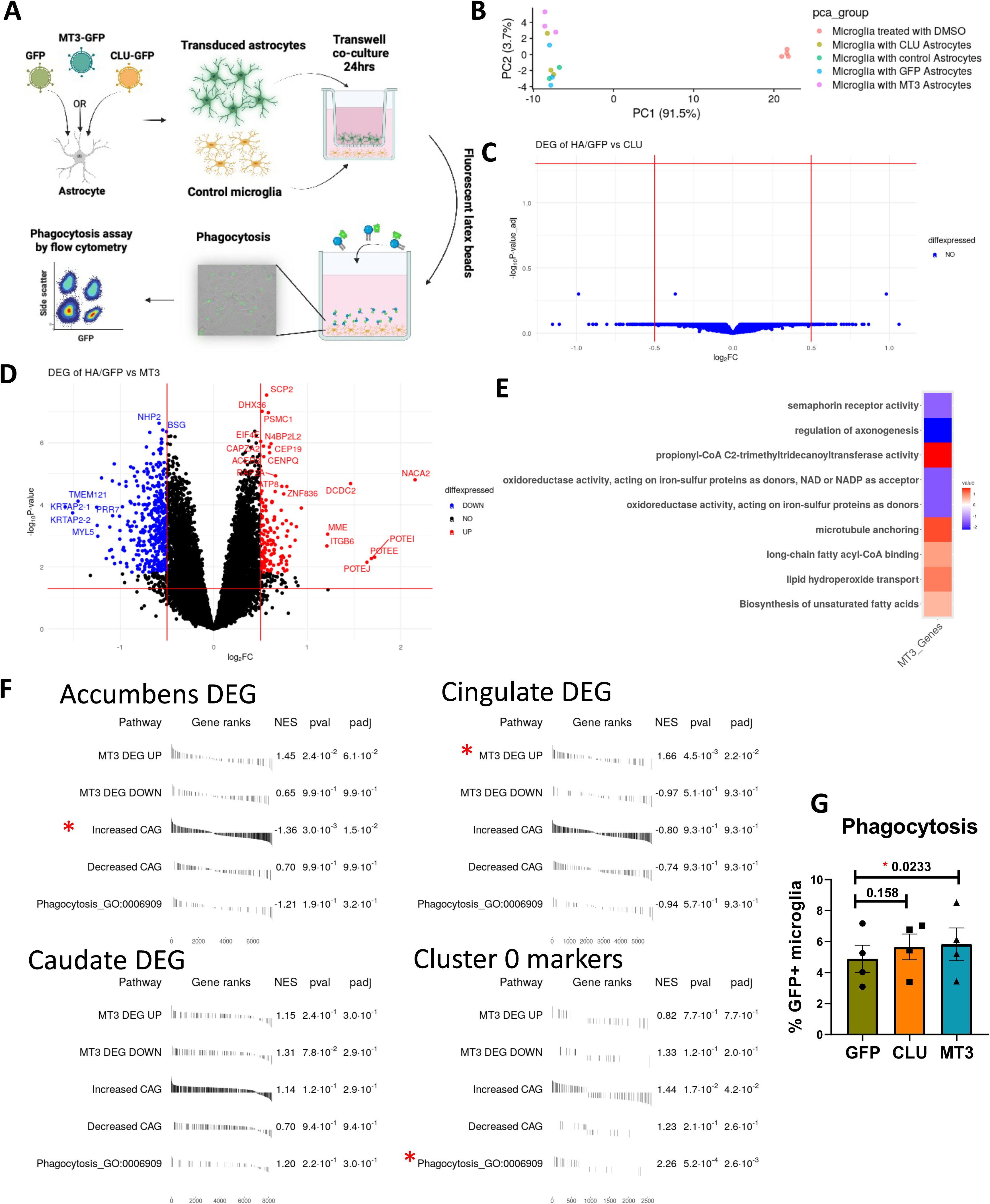
MT3 astrocytes modulate microglial function. **A)** A cartoon schema depicting the trans-well co-culture experiment between astrocytes and microglia (HMC3 cells). **B)** PCA plot of the first two component of the bulk-RNAseq samples: control samples are co-cultures with GFP astrocytes or untransduced astrocytes, experimental samples include microglia co-cultured with CLU or MT3 overexpressing astrocytes. Mono-cultures of microglial cells are also shown. Note that co-culture is a major driver of gene expression. **C)** Volcano plot showing that no genes were differentially expressed between the control (HA/GFP = human astrocytes or GFP transduced astrocyte control group co-cultures) versus microglia with CLU astrocytes. **D)** Volcano plot of the differentially expressed comparing the same control in (**C**) versus microglia co-cultured with MT3 astrocytes (genes in red are significantly increased in the MT3 group and genes in blue are significantly decreased in MT3). **E)** Heatmap displaying the log10(p-value) of the GO terms of the DEG from (**D**). **F)** Pre-ranked gene set enrichment analysis determining the enrichment of DEG of MT3 from **(D),** CAG-correlated gene sets from **Figure 1G**, and a GO phagocytosis term gene, in the ranked DEG from **Figure S10C-E** and ranked cluster 0 markers (cluster 0 = microglial cluster from **Figure S10A**). The adjusted p values and normalized enrichment scores (NES) are indicated. Red stars indicate significance. **G)** Bar plot showing the percentage of microglia with GFP beads that have been phagocytized in three scenarios (GFP = Microglia in co-culture with GFP astrocytes, CLU = Microglia in co-culture with CLU astrocytes, MT3 = Microglia in co-culture with MT3 astrocytes). N = 4 biological replicates. Paired two-tailed t-tests. The p values are indicated.

## References

1. Hirano, M. et al. Clinicopathological differences between the motor onset and psychiatric onset of Huntington’s disease, focusing on the nucleus accumbens. Neuropathology 39, 331–341 (2019).

2. Hebb, M.O., Denovan-Wright, E.M. & Robertson, H.A. Expression of the Huntington’s disease gene is regulated in astrocytes in the arcuate nucleus of the hypothalamus of postpartum rats. FASEB J 13, 1099–1106 (1999).

3. DiFiglia, M. et al. Aggregation of huntingtin in neuronal intranuclear inclusions and dystrophic neurites in brain. Science 277, 1990–1993 (1997).

4. Mangiarini, L. et al. Exon 1 of the HD gene with an expanded CAG repeat is sufficient to cause a progressive neurological phenotype in transgenic mice. Cell 87, 493–506 (1996).

5. Vonsattel, J.P. et al. Neuropathological classification of Huntington’s disease. J Neuropathol Exp Neurol 44, 559–577 (1985).

6. Roos, R.A., Pruyt, J.F., de Vries, J. & Bots, G.T. Neuronal distribution in the putamen in Huntington’s disease. J Neurol Neurosurg Psychiatry 48, 422–425 (1985).

7. Rosas, H.D. et al. Cerebral cortex and the clinical expression of Huntington’s disease: complexity and heterogeneity. Brain 131, 1057–1068 (2008).

8. Iennaco, R. et al. The evolutionary history of the polyQ tract in huntingtin sheds light on its functional pro-neural activities. Cell death and differentiation 29, 293–305 (2022).

9. Roy, J.C.L. et al. Somatic CAG expansion in Huntington’s disease is dependent on the MLH3 endonuclease domain, which can be excluded via splice redirection. Nucleic Acids Res 49, 3907–3918 (2021).

10. Gonitel, R. et al. DNA instability in postmitotic neurons. Proceedings of the National Academy of Sciences of the United States of America 105, 3467–3472 (2008).

11. Shelbourne, P.F. et al. Triplet repeat mutation length gains correlate with cell-type specific vulnerability in Huntington disease brain. Human molecular genetics 16, 1133–1142 (2007).

12. Wang, N. et al. Neuronal targets for reducing mutant huntingtin expression to ameliorate disease in a mouse model of Huntington’s disease. Nature medicine 20, 536–541 (2014).

13. Swami, M. et al. Somatic expansion of the Huntington’s disease CAG repeat in the brain is associated with an earlier age of disease onset. Human molecular genetics 18, 3039–3047 (2009).

14. Zuccato, C. et al. Loss of huntingtin-mediated BDNF gene transcription in Huntington’s disease. Science 293, 493–498 (2001).

15. Lim, R.G. et al. Huntington disease oligodendrocyte maturation deficits revealed by single-nucleus RNAseq are rescued by thiamine-biotin supplementation. Nature communications 13, 7791 (2022).

16. Vonsattel, J.P., Keller, C. & Del Pilar Amaya, M. Neuropathology of Huntington’s disease. Handb Clin Neurol 89, 599–618 (2008).

17. Jiang, R., Diaz-Castro, B., Looger, L.L. & Khakh, B.S. Dysfunctional Calcium and Glutamate Signaling in Striatal Astrocytes from Huntington’s Disease Model Mice. The Journal of neuroscience : the official journal of the Society for Neuroscience 36, 3453–3470 (2016).

18. Faideau, M. et al. In vivo expression of polyglutamine-expanded huntingtin by mouse striatal astrocytes impairs glutamate transport: a correlation with Huntington’s disease subjects. Human molecular genetics 19, 3053–3067 (2010).

19. Lievens, J.C. et al. Impaired glutamate uptake in the R6 Huntington’s disease transgenic mice. Neurobiology of disease 8, 807–821 (2001).

20. Diaz-Castro, B., Gangwani, M.R., Yu, X., Coppola, G. & Khakh, B.S. Astrocyte molecular signatures in Huntington’s disease. Sci Transl Med 11 (2019).

21. Al-Dalahmah, O. et al. Single-nucleus RNA-seq identifies Huntington disease astrocyte states. Acta Neuropathol Commun 8, 19 (2020).

22. Bradford, J. et al. Mutant huntingtin in glial cells exacerbates neurological symptoms of Huntington disease mice. The Journal of biological chemistry 285, 10653–10661 (2010).

23. Bradford, J. et al. Expression of mutant huntingtin in mouse brain astrocytes causes age- dependent neurological symptoms. Proceedings of the National Academy of Sciences of the United States of America 106, 22480–22485 (2009).

24. Hsiao, H.Y., Chen, Y.C., Chen, H.M., Tu, P.H. & Chern, Y. A critical role of astrocyte-mediated nuclear factor-kappaB-dependent inflammation in Huntington’s disease. Human molecular genetics 22, 1826–1842 (2013).

25. Wood, T.E. et al. Mutant huntingtin reduction in astrocytes slows disease progression in the BACHD conditional Huntington’s disease mouse model. Human molecular genetics 28, 487–500 (2019).

26. Yu, X. et al. Reducing Astrocyte Calcium Signaling In Vivo Alters Striatal Microcircuits and Causes Repetitive Behavior. Neuron 99, 1170–1187 e1179 (2018).

27. Benraiss, A. et al. Cell-intrinsic glial pathology is conserved across human and murine models of Huntington’s disease. Cell reports 36, 109308 (2021).

28. Reyes-Ortiz, A.M. et al. Single-nuclei transcriptome analysis of Huntington disease iPSC and mouse astrocytes implicates maturation and functional deficits. iScience 26, 105732 (2023).

29. Meunier, C., Merienne, N., Jolle, C., Deglon, N. & Pellerin, L. Astrocytes are key but indirect contributors to the development of the symptomatology and pathophysiology of Huntington’s disease. Glia 64, 1841–1856 (2016).

30. Benraiss, A. et al. Human glia can both induce and rescue aspects of disease phenotype in Huntington disease. Nature communications 7, 11758 (2016).

31. Tong, X. et al. Astrocyte Kir4.1 ion channel deficits contribute to neuronal dysfunction in Huntington’s disease model mice. Nature neuroscience 17, 694–703 (2014).

32. Arregui, L., Benitez, J.A., Razgado, L.F., Vergara, P. & Segovia, J. Adenoviral astrocyte-specific expression of BDNF in the striata of mice transgenic for Huntington’s disease delays the onset of the motor phenotype. Cellular and molecular neurobiology 31, 1229–1243 (2011).

33. Giralt, A. et al. BDNF regulation under GFAP promoter provides engineered astrocytes as a new approach for long-term protection in Huntington’s disease. Gene therapy 17, 1294–1308 (2010).

34. Oliveira, A.O., Osmand, A., Outeiro, T.F., Muchowski, P.J. & Finkbeiner, S. alphaB-Crystallin overexpression in astrocytes modulates the phenotype of the BACHD mouse model of Huntington’s disease. Human molecular genetics 25, 1677–1689 (2016).

35. Rüb, U., Vonsattell, J.P.G., Heinsen, H. & Korf, H.-W. The Neuropathology of Huntington’s Disease: Classical Findings, Recent Developments and Correlation to Functional Neuroanatomy. (2015).

36. Labadorf, A. et al. RNA Sequence Analysis of Human Huntington Disease Brain Reveals an Extensive Increase in Inflammatory and Developmental Gene Expression, in PLoS One, Vol. 10 (2015).

37. Hodges, A. et al. Regional and cellular gene expression changes in human Huntington’s disease brain. Human molecular genetics 15, 965–977 (2006).

38. Garcia, F.J. et al. Single-cell dissection of the human brain vasculature. Nature 603, 893–899 (2022).

39. Lee, H. et al. Cell Type-Specific Transcriptomics Reveals that Mutant Huntingtin Leads to Mitochondrial RNA Release and Neuronal Innate Immune Activation. Neuron (2020).

40. Tyagi, E., Fiorelli, T., Norden, M. & Padmanabhan, J. Alpha 1-Antichymotrypsin, an Inflammatory Protein Overexpressed in the Brains of Patients with Alzheimer’s Disease, Induces Tau Hyperphosphorylation through c-Jun N-Terminal Kinase Activation. Int J Alzheimers Dis 2013, 606083 (2013).

41. Block, R.C., Dorsey, E.R., Beck, C.A., Brenna, J.T. & Shoulson, I. Altered cholesterol and fatty acid metabolism in Huntington disease. J Clin Lipidol 4, 17–23 (2010).

42. Kreilaus, F., Spiro, A.S., Hannan, A.J., Garner, B. & Jenner, A.M. Brain Cholesterol Synthesis and Metabolism is Progressively Disturbed in the R6/1 Mouse Model of Huntington’s Disease: A Targeted GC-MS/MS Sterol Analysis. J Huntingtons Dis 4, 305–318 (2015).

43. Kreilaus, F., Spiro, A.S., McLean, C.A., Garner, B. & Jenner, A.M. Evidence for altered cholesterol metabolism in Huntington’s disease post mortem brain tissue. Neuropathology and applied neurobiology 42, 535–546 (2016).

44. Phillips, G.R. et al. Cholesteryl ester levels are elevated in the caudate and putamen of Huntington’s disease patients. Sci Rep 10, 20314 (2020).

45. Valenza, M. et al. Cholesterol biosynthesis pathway is disturbed in YAC128 mice and is modulated by huntingtin mutation. Human molecular genetics 16, 2187–2198 (2007).

46. Valenza, M. et al. Progressive dysfunction of the cholesterol biosynthesis pathway in the R6/2 mouse model of Huntington’s disease. Neurobiology of disease 28, 133–142 (2007).

47. Valenza, M. et al. Dysfunction of the cholesterol biosynthetic pathway in Huntington’s disease. The Journal of neuroscience : the official journal of the Society for Neuroscience 25, 9932–9939 (2005).

48. Lee, J.A., Hall, B., Allsop, J., Alqarni, R. & Allen, S.P. Lipid metabolism in astrocytic structure and function. Seminars in cell & developmental biology 112, 123–136 (2021).

49. Al Dalahmah, O., et al. The Matrix Receptor CD44 Is Present in Astrocytes Throughout the Human CNS and Accumulates in Hypoxia and Seizures. Preprints.org (2023).

50. Sosunov, A.A. et al. Phenotypic heterogeneity and plasticity of isocortical and hippocampal astrocytes in the human brain. The Journal of neuroscience : the official journal of the Society for Neuroscience 34, 2285–2298 (2014).

51. (!!! INVALID CITATION !!! 21).

52. Isani, G. & Carpene, E. Metallothioneins, unconventional proteins from unconventional animals: a long journey from nematodes to mammals. Biomolecules 4, 435–457 (2014).

53. Kuchroo, M. et al. Multiscale PHATE identifies multimodal signatures of COVID-19. Nat Biotechnol 40, 681–691 (2022).

54. Farmer, B.C., Kluemper, J. & Johnson, L.A. Apolipoprotein E4 Alters Astrocyte Fatty Acid Metabolism and Lipid Droplet Formation. Cells 8 (2019).

55. Lovatt, D. et al. The transcriptome and metabolic gene signature of protoplasmic astrocytes in the adult murine cortex. The Journal of neuroscience : the official journal of the Society for Neuroscience 27, 12255–12266 (2007).

56. Xie, Z. et al. Gene Set Knowledge Discovery with Enrichr. Curr Protoc 1, e90 (2021).

57. Lachmann, A. et al. ChEA: transcription factor regulation inferred from integrating genome-wide ChIP-X experiments. Bioinformatics 26, 2438–2444 (2010).

58. Glass, M., Dragunow, M. & Faull, R.L. The pattern of neurodegeneration in Huntington’s disease: a comparative study of cannabinoid, dopamine, adenosine and GABA(A) receptor alterations in the human basal ganglia in Huntington’s disease. Neuroscience 97, 505–519 (2000).

59. Reiner, A. et al. Differential loss of striatal projection neurons in Huntington disease. Proceedings of the National Academy of Sciences of the United States of America 85, 5733–5737 (1988).

60. Sieradzan, K.A. & Mann, D.M. The selective vulnerability of nerve cells in Huntington’s disease. Neuropathology and applied neurobiology 27, 1–21 (2001).

61. Deng, Y.P. et al. Differential loss of striatal projection systems in Huntington’s disease: a quantitative immunohistochemical study. Journal of chemical neuroanatomy 27, 143–164 (2004).

62. Matsushima, A. et al. Transcriptional vulnerabilities of striatal neurons in human and rodent models of Huntington’s disease. Nature communications 14, 282 (2023).

63. Hedreen, J.C. & Folstein, S.E. Early loss of neostriatal striosome neurons in Huntington’s disease. J Neuropathol Exp Neurol 54, 105–120 (1995).

64. Morton, A.J., Nicholson, L.F. & Faull, R.L. Compartmental loss of NADPH diaphorase in the neuropil of the human striatum in Huntington’s disease. Neuroscience 53, 159–168 (1993).

65. Wojtas, A.M. et al. Astrocyte-derived clusterin suppresses amyloid formation in vivo. Molecular neurodegeneration 15, 71 (2020).

66. Chen, F. et al. Clusterin secreted from astrocyte promotes excitatory synaptic transmission and ameliorates Alzheimer’s disease neuropathology. Molecular neurodegeneration 16, 5 (2021).

67. Querol-Vilaseca, M. et al. YKL-40 (Chitinase 3-like I) is expressed in a subset of astrocytes in Alzheimer’s disease and other tauopathies. Journal of neuroinflammation 14, 118 (2017).

68. Lananna, B.V. et al. Chi3l1/YKL-40 is controlled by the astrocyte circadian clock and regulates neuroinflammation and Alzheimer’s disease pathogenesis. Science Translational Medicine 12 (2020).

69. Madden, N. et al. The link between SARS-CoV-2 related microglial reactivity and astrocyte pathology in the inferior olivary nucleus. Frontiers in neuroscience 17 (2023).

70. Donaldson, J., Powell, S., Rickards, N., Holmans, P. & Jones, L. What is the Pathogenic CAG Expansion Length in Huntington’s Disease? J Huntingtons Dis 10, 175–202 (2021).

71. (!!! INVALID CITATION !!! 61-63).

72. Ng, B. et al. An xQTL map integrates the genetic architecture of the human brain’s transcriptome and epigenome. Nature neuroscience 20, 1418–1426 (2017).

73. Murakami, S., Miyazaki, I., Sogawa, N., Miyoshi, K. & Asanuma, M. Neuroprotective effects of metallothionein against rotenone-induced myenteric neurodegeneration in parkinsonian mice. Neurotox Res 26, 285–298 (2014).

74. Victor, M.B. et al. Striatal neurons directly converted from Huntington’s disease patient fibroblasts recapitulate age-associated disease phenotypes. Nature neuroscience 21, 341–352 (2018).

75. Kalathur, R.K. et al. The unfolded protein response and its potential role in Huntington’s disease elucidated by a systems biology approach. F1000Res 4, 103 (2015).

76. Trendelenburg, G. et al. Serial analysis of gene expression identifies metallothionein-II as major neuroprotective gene in mouse focal cerebral ischemia. The Journal of neuroscience : the official journal of the Society for Neuroscience 22, 5879–5888 (2002).

77. Chung, R.S. et al. Neuron-glia communication: metallothionein expression is specifically up- regulated by astrocytes in response to neuronal injury. J Neurochem 88, 454–461 (2004).

78. Stankovic, R.K., Chung, R.S. & Penkowa, M. Metallothioneins I and II: neuroprotective significance during CNS pathology. The international journal of biochemistry & cell biology 39, 484–489 (2007).

79. Michael, G.J. et al. Up-regulation of metallothionein gene expression in parkinsonian astrocytes. Neurogenetics 12, 295–305 (2011).

80. Uchida, Y., Gomi, F., Masumizu, T. & Miura, Y. Growth inhibitory factor prevents neurite extension and the death of cortical neurons caused by high oxygen exposure through hydroxyl radical scavenging. The Journal of biological chemistry 277, 32353–32359 (2002).

81. Erickson, J.C., Hollopeter, G., Thomas, S.A., Froelick, G.J. & Palmiter, R.D. Disruption of the metallothionein-III gene in mice: analysis of brain zinc, behavior, and neuron vulnerability to metals, aging, and seizures. The Journal of neuroscience : the official journal of the Society for Neuroscience 17, 1271–1281 (1997).

82. Koh, J.Y. & Lee, S.J. Metallothionein-3 as a multifunctional player in the control of cellular processes and diseases. Molecular brain 13, 116 (2020).

83. Chung, R.S. et al. Redefining the role of metallothionein within the injured brain: extracellular metallothioneins play an important role in the astrocyte-neuron response to injury. The Journal of biological chemistry 283, 15349–15358 (2008).

84. Klaassen, C.D., Liu, J. & Diwan, B.A. Metallothionein protection of cadmium toxicity. Toxicol Appl Pharmacol 238, 215–220 (2009).

85. Rivera-Mancia, S. et al. The transition metals copper and iron in neurodegenerative diseases. Chemico-biological interactions 186, 184–199 (2010).

86. Choi, D.W. Glutamate neurotoxicity and diseases of the nervous system. Neuron 1, 623–634 (1988).

87. Kohyama, J. et al. BMP-induced REST regulates the establishment and maintenance of astrocytic identity. The Journal of cell biology 189, 159–170 (2010).

88. Hirabayashi, Y. et al. Polycomb limits the neurogenic competence of neural precursor cells to promote astrogenic fate transition. Neuron 63, 600–613 (2009).

89. Dobin, A. et al. STAR: ultrafast universal RNA-seq aligner. Bioinformatics 29, 15–21 (2013).

90. Robinson, M.D., McCarthy, D.J. & Smyth, G.K. edgeR: a Bioconductor package for differential expression analysis of digital gene expression data. Bioinformatics 26, 139–140 (2010).

91. Kuleshov, M.V. et al. Enrichr: a comprehensive gene set enrichment analysis web server 2016 update. Nucleic Acids Res 44, W90–97 (2016).

92. Liis Kolberg & Raudvere, U., Edn. 0.2.0. (2020).

93. Raudvere, U. et al. g:Profiler: a web server for functional enrichment analysis and conversions of gene lists (2019 update). Nucleic Acids Res 47, W191–W198 (2019).

94. Bligh, E.G. & Dyer, W.J. A rapid method of total lipid extraction and purification. Can J Biochem Physiol 37, 911–917 (1959).

95. Chan, R.B. et al. Comparative lipidomic analysis of mouse and human brain with Alzheimer disease. J Biol Chem 287, 2678–2688 (2012).

96. Hsu, F.F., Turk, J., Shi, Y. & Groisman, E.A. Characterization of acylphosphatidylglycerols from Salmonella typhimurium by tandem mass spectrometry with electrospray ionization. J Am Soc Mass Spectrom 15, 1–11 (2004).

97. Langfelder, P. & Horvath, S. WGCNA: an R package for weighted correlation network analysis. BMC Bioinformatics 9, 559 (2008).

98. Ritchie, M.E. et al. limma powers differential expression analyses for RNA-sequencing and microarray studies. Nucleic Acids Res 43, e47 (2015).

99. Johnson, W.E., Li, C. & Rabinovic, A. Adjusting batch effects in microarray expression data using empirical Bayes methods. Biostatistics 8, 118–127 (2007).

100. Langfelder, P., Luo, R., Oldham, M.C. & Horvath, S. Is my network module preserved and reproducible? PLoS Comput Biol 7, e1001057 (2011).

101. Rohart, F., Gautier, B., Singh, A. & Le Cao, K.A. mixOmics: An R package for ‘omics feature selection and multiple data integration. PLoS Comput Biol 13, e1005752 (2017).

102. McCarthy, D.J., Campbell, K.R., Lun, A.T. & Wills, Q.F. Scater: pre-processing, quality control, normalization and visualization of single-cell RNA-seq data in R. Bioinformatics 33, 1179–1186 (2017).

103. Shannon, P. et al. Cytoscape: a software environment for integrated models of biomolecular interaction networks. Genome research 13, 2498–2504 (2003).

104. Street, K. et al. Slingshot: cell lineage and pseudotime inference for single-cell transcriptomics. BMC Genomics 19, 477 (2018).

105. Van den Berge, K., et al. Trajectory-based differential expression analysis for single-cell sequencing data. Nature communications 11, 1201 (2020).

106. Lin, H. & Peddada, S.D. Analysis of compositions of microbiomes with bias correction. Nature communications 11, 3514 (2020).

107. Genetic Modifiers of Huntington’s Disease, C. Identification of Genetic Factors that Modify Clinical Onset of Huntington’s Disease. Cell 162, 516–526 (2015).

108. Oh, Y.M. et al. Age-related Huntington’s disease progression modeled in directly reprogrammed patient-derived striatal neurons highlights impaired autophagy. Nature neuroscience 25, 1420–1433 (2022).

## References

1. Chen, T., Gai, W.P. & Abbott, C.A. Dipeptidyl peptidase 10 (DPP10(789)): a voltage gated potassium channel associated protein is abnormally expressed in Alzheimer’s and other neurodegenerative diseases. Biomed Res Int 2014, 209398 (2014).

2. Runne, H. et al. Dysregulation of gene expression in primary neuron models of Huntington’s disease shows that polyglutamine-related effects on the striatal transcriptome may not be dependent on brain circuitry. The Journal of neuroscience : the official journal of the Society for Neuroscience 28, 9723–9731 (2008).

3. Niccolini, F. et al. Altered PDE10A expression detectable early before symptomatic onset in Huntington’s disease. Brain 138, 3016–3029 (2015).

4. Malaiya, S. et al. Single-Nucleus RNA-Seq Reveals Dysregulation of Striatal Cell Identity Due to Huntington’s Disease Mutations. The Journal of neuroscience : the official journal of the Society for Neuroscience 41, 5534–5552 (2021).

5. Lee, H. et al. Cell Type-Specific Transcriptomics Reveals that Mutant Huntingtin Leads to Mitochondrial RNA Release and Neuronal Innate Immune Activation. Neuron (2020).

6. Matejuk, A. & Ransohoff, R.M. Crosstalk Between Astrocytes and Microglia: An Overview. Front Immunol 11, 1416 (2020).

7. Thrupp, N. et al. Single-Nucleus RNA-Seq Is Not Suitable for Detection of Microglial Activation Genes in Humans. Cell reports 32, 108189 (2020).

8. Taban, Q., Mumtaz, P.T., Masoodi, K.Z., Haq, E. & Ahmad, S.M. Scavenger receptors in host defense: from functional aspects to mode of action. Cell Commun Signal 20, 2 (2022).

9. Al-Dalahmah, O. et al. Re-convolving the compositional landscape of primary and recurrent glioblastoma reveals prognostic and targetable tissue states. Nature communications 14, 2586 (2023).

